# Spatial transcriptomics identifies distinct domains regulating yield-related traits of the wheat ear

**DOI:** 10.1101/2025.08.12.670006

**Authors:** Yue Qu, Cong Tan, Liujing Yang, Marianna Pasquariello, Abdul Kader Alabdullah, Shiyu Sun, Munir Iqbal, John Salamon, Scott Boden

**Affiliations:** School of Agriculture, Food and Wine, Waite Research Institute, University of Adelaide, Glen Osmond, SA 5064, Australia; Key Laboratory of Genomics, Ministry of Agricultural, BGI Bioverse, BGI Research, Shenzhen, 518083, China; College of Life Sciences, University of Chinese Academy of Sciences, Beijing 100049, China; Crop Genetics Department, John Innes Centre, Norwich Research Park, Norwich, NR4 7UH, UK; Council for Agricultural Research and Economics, Research Centre for Genomics and Bioinformatics, Via San Protaso 69, Fiorenzuola d’ Arda (PC), 29017, Italy; South Australian Genomics Centre, SAHMRI, North Terrace, Adelaide, SA 5000, Australia

**Author notes:** These authors contributed equally: Yue Qu, Cong Tan.

## Abstract

Cereal inflorescences are complex, highly ordered structures composed of grain-producing florets that form within specialised branches called spikelets. The spikelets of wheat are arranged in two alternating rows along a central rachis, in a pattern determined during early reproductive development. While several genes that control spikelet development have been identified, the molecular processes that regulate their morphology and the formation of supporting structures, such as meristems and the rachis, remain poorly understood. Here, we used spatial transcriptomics to investigate the dynamic transcriptional landscape of a wheat inflorescence during spikelet development. We identified two spatially distinct regions that regulate spikelet architecture, including a primordium region characterised by *RAMOSA2* activity, and a boundary region that expresses *ALOG1* and known regulators of bract suppression. Developmental assays indicate that spikelets differentiate from meristematic regions, which is accompanied by formation of central vascular regions of the rachis and inflorescence base that express genes controlling spikelet number. The combined spatial transcriptome and genetic data reveal key regulators of spikelet development, including target genes for improving spikelet number and yield.

## INTRODUCTION

The inflorescence of bread wheat (*Triticum aestivum*), known as the spike, is a complex and highly ordered structure comprised of spikelets, which form grain-bearing florets. As grain number is a key yield determinant, the number of spikelets and fertile florets that form on a spike is crucial for global food security, especially because wheat accounts for 20% of the calories and protein consumed worldwide^1^. The early inflorescence stages of double ridge (DR) and lemma primordium (LP) mark a pivotal developmental phase that establishes the foundation for the final spike architecture and potential grain number^2,3^. During the DR stage, the inflorescence meristem differentiates to generate distinct lateral meristems destined to become spikelets, marking the transition from vegetative to reproductive growth^4^. At the LP stage, individual spikelets differentiate further to initiate lemma primordia and floret meristems, setting the groundwork for floret formation and grain production^4,5^.

Given their importance for grain production, many studies have sought to identify genes that regulate the number, arrangement, and fertility of spikelets that form on the wheat spike. Genes including *Photoperiod-1* (*Ppd-1*)^6^, *FLOWERING LOCUS T2* (*FT2*)^7^ and *PHOTOPERIOD-1 DEPENDENT bZIP TRANSCRIPTION FACTOR1* (*PDB1*)^3^, *CONSTANS-LIKE5* (*COL5*)^8^, *ZINC FINGER1* (*ZF1*)^9^ and *TERMINAL FLOWER1* (*TFL1*)^2^ influence spikelet number by modulating the strength of the flowering signal or the timing of spikelet termination, with many expressed during DR or LP. Spikelet architecture genes–such as *ALOG1*^3^*, DUO1*^10^*, TEOSINTE BRANCHED1* (*TB1)*^11^*, HOMEOBOX DOMAIN-2* (*HB-2*)^12^, and *WHEAT FRIZZY PANCILE* (*WFZP*)^13,14^–have been identified by analysing genotypes that form supernumerary spikelets or branched spikes, and they are similarly expressed during these early developmental stages. Spikelet fertility is largely regulated by MADS-box and homeodomain leucine zipper class I transcription factors, SHORT VEGETATIVE PHASE1/2 (SVP1/2)^15–18^ and GRAIN NUMBER INCREASE1 (GNI1)^19^, which influence either the fertility of basal spikelets or the survival of florets within each spikelet. While these studies highlight the emerging capabilities of wheat developmental genetics, it is sobering that each gene was identified individually through lengthy studies reliant on unique natural or induced variant alleles, that are often dominant due to the redundancy of wheat’s hexaploid genome. This shortcoming highlights the need for new approaches that enable broader and more rapid assessment of genes influencing wheat inflorescence development.

Spatial transcriptomics is an emerging and rapidly advancing technology that enables unprecedented spatial resolution of gene expression within intact tissue structures, overcoming limitations of conventional transcriptome approaches that analyse bulk tissue samples^3,20,21^. Recent spatial transcriptomics studies have demonstrated the potential of this approach to enhance our understanding of cereal development; in maize, spatial transcriptomics identified distinct meristem subtypes during ear development and uncovered spatially-specific expression of key regulatory genes^22^. Similarly, spatial transcriptomic analyses in barley have demonstrated dynamic gene expression during seed germination, revealing intricate tissue-specific regulatory patterns^23^. These advances highlight spatial transcriptomics’ potential to elucidate previously inaccessible gene expression patterns that control early spike development, especially in wheat where next generation sequencing can resolve homeologous transcripts of the three genomes (A, B and D). Clarifying these expression patterns could help bridge existing knowledge gaps about spikelet formation, ultimately informing targeted wheat breeding strategies designed to optimize inflorescence architecture and enhance grain yield potential.

In this study, we used Stereo-seq (SpaTial Enhanced Resolution Omics sequencing) to generate a spatially resolved, single-cell transcriptome atlas of the wheat inflorescence at the DR and LP developmental stages. This high-resolution approach revealed novel spatial patterns of gene expression associated with spikelet initiation and differentiation. By identifying key regulatory regions that control spikelet number and arrangement, our findings provide a foundation for targeted strategies to improve wheat yield.

## RESULTS

### High-resolution spatial transcriptome profiling of wheat inflorescences using Stereo-seq

To investigate cell differentiation and cell fate determine that associate with spikelet development in wheat, we performed spatial transcriptomic profiling for inflorescences at the double ridge and lemma primordium stages that mark the beginning and completion of spikelet initiation^7^. Developing spikes from the elite cultivar Mace were flash-frozen in optimal cutting temperature resin, sectioned longitudinally at 12 µm, and mounted onto 1 cm x 1 cm STOMics Stereo-seq chips. These sections were subjected to a workflow that included nuclei and cell wall staining, analysis of tissue integrity, tissue permeabilization, cDNA synthesis, library generation and sequencing, in preparation for the downstream analyses (Fig. 1a). We obtained high quality spatial transcriptomes (1.76 billion high-confidence reads) for four sections of the double ridge and lemma primordium stages, with means of 48,178 and 53,373 genes detected across the four replicate sections for DR and LP, respectively, which represents 89.6% and 96.3% of expressed transcripts detected using bulk RNA-seq analysis of the same stages^3^. The high-resolution stained images and sequencing data of the four sections for each stage were used to perform spatial clustering at different bin sizes (Fig. 1b,c; Supplementary Table 1 and Supplementary Figure 1,2). The bin40 and bin50, which correspond to approximately one cell of a developing wheat inflorescence, defined regions of the inflorescence that represented structural features (e.g., spikelet primordia, rachis, inflorescence meristem, lateral meristems) for the DR and LP stages, respectively, and these bin sizes were used for further analyses.

**Figure 1.**
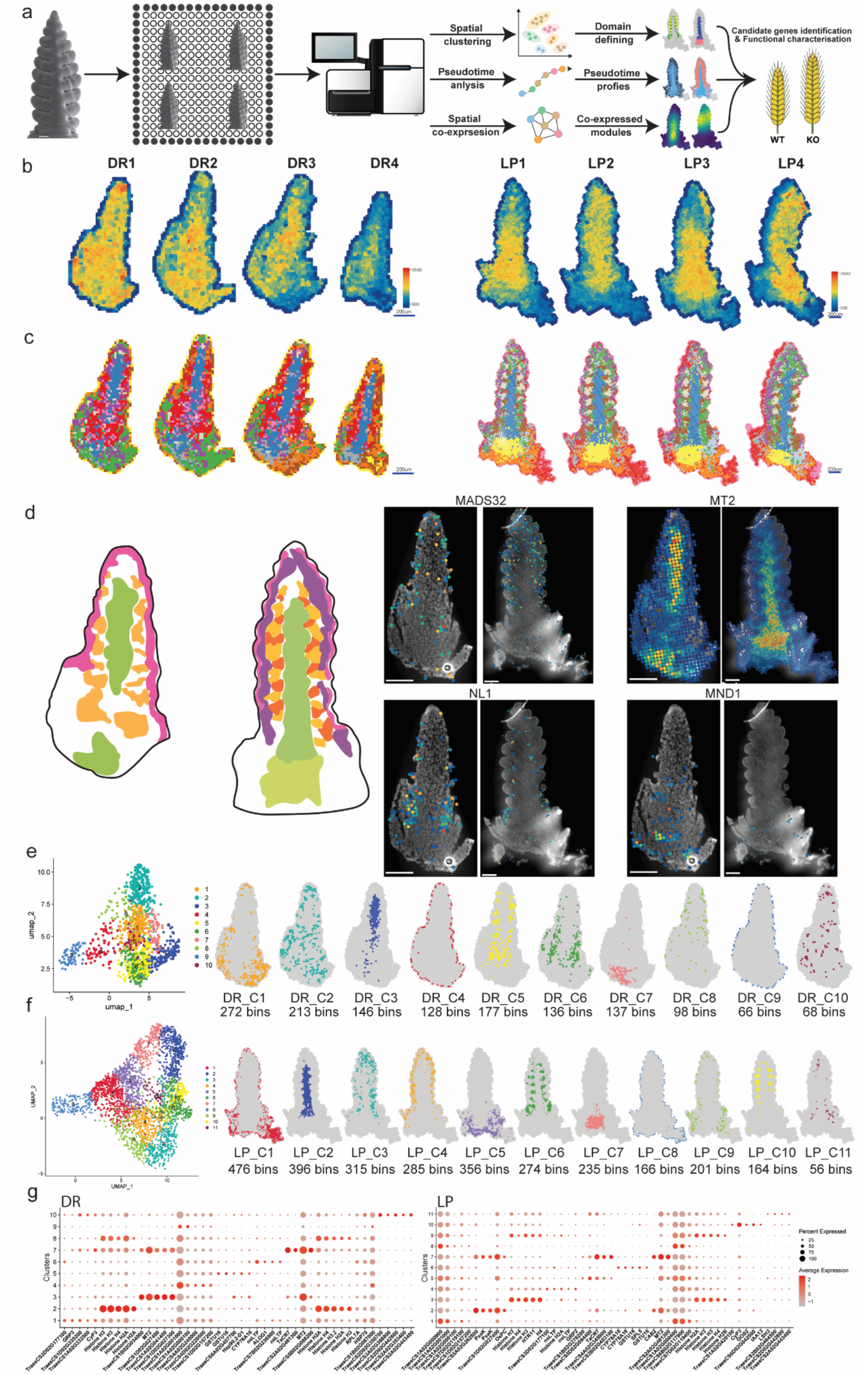
Spatial transcriptomics workflow and identification of transcriptional domains in early wheat inflorescence development. **a**, Schematic overview of the spatial transcriptomics pipeline that informs candidate gene identification and functional validation. **b**, Spatial transcript count maps and **c**, spatial binning resolution optimization across four biological replicates of double ridge (DR1–DR4) and lemma primordium (LP1–LP4) stages. Heatmaps indicate unique molecular identifier (UMI) counts per bin, and representative spatial maps of DR (bin40) and LP (bin50) show selected bin sizes based on a balance between spatial resolution and transcript capture (see Supplementary Table 1). Scale bars, 200 µm. **d**, Schematic of spatial gene expression patterns that overlap with key inflorescence tissues: meristems (pink and purple), rachis and basal vasculature (dark and light green) and spikelets (light and dark orange). Gene expression patterns of *MADS32*, *MT2*, *NL1*, and *MND1* are localized to distinct regions of the inflorescence, which overlap with previous *in situ* hybridization or MERFISH data (Supplementary Figure 2). Scale bars, 200 µm. **e**, Unsupervised clustering of DR transcriptomes. UMAP projection of spatial bins reveals 10 transcriptionally distinct clusters (DR_C1–DR_C10). **f**, Unsupervised clustering of LP transcriptomes. UMAP projection shows 11 clusters (LP_C1–LP_C11) capturing domain-specific expression in the LP stage. Spatial maps depict localization of individual clusters. Bin numbers are indicated for each cluster (**e, f**). **g**, Top cluster-specific genes. Dot plots displaying the five most enriched genes per cluster for both DR and LP. Dot size represents the proportion of bins expressing each gene; color indicates average expression level.

We validated the spatial gene expression data by comparing our results with those obtained in other studies using independent assays. Homeologous transcripts for key marker genes including *NECK LEAF1* (*NL1*), *MADS32*, *METALLOTHIONEIN2* (*MT2*), *MANY NODED DWARF1* (*MND1*), *ALOG1*, *LAX PANICLE1* (*LAX1*), *VEGETATIVE TO REPRODUCTIVE TRANSITION2* (*VRT2*) and etc., exhibited spatially distinct expression patterns consistent with known tissue-specific localization during inflorescence development^3,24–26^ (Fig. 1d; Supplementary Figure 3-12). For example, *NL1* and *ALOG1* transcripts localised to the lower region of the lateral meristem that subtends spikelet primordia, while *MADS32* and *LAX1* transcripts localised to the initiating spikelet primordia and the base of differentiated spikelets at DR and LP, respectively. *MT2* transcripts localised to the central vasculature of the developing rachis at both stages, matching the phloem specific expression of rice orthologues, and *MND1* transcripts were detected in vasculature at the inflorescence base, with a stronger signal detected at DR than LP. Our data, therefore, align closely with spatial patterns of gene expression detected using complementary approaches across diverse inflorescence tissues^3,24–26^.

### Spatial clustering defines distinct transcriptional domains during early inflorescence development

To investigate transcriptional domains in the developing inflorescence, we performed unsupervised clustering analysis using all four sections for each of the DR and LP stages. This analysis detected 10 clusters at DR and 11 at LP, which resolved into spatially distinct regions based on their uniform manifold approximation and projection (UMAP) (Fig. 1e,f). Two DR clusters, designated the spikelet initiation (DR_C5) and leaf ridge (DR_C6) clusters, located to regions lateral to the main axis where spikelet and leaf primordia form, respectively. Similarly, two distinct LP clusters, termed spikelet primordia (LP_C10) and spikelet boundary (LP_C6), were arranged distichously along the central axis that aligned with developing spikelets. We also detected domains at both stages that were characteristic of the central rachis and basal vasculature of the inflorescence (DR_C3, DR_C7; LP_C2, LP_C7, respectively), and others localised to the apical inflorescence and lateral meristem regions (DR_C1, DR_C2, DR_C8 and DR_C10; LP_C3, LP_C9, LP_C4 and LP_C11, respectively). The clusters were defined by marker genes that exhibited strong expression in the representative bins (Fig. 1g; Supplementary Table 2). The spikelet initiation cluster at DR (DR_C5) was defined by homeologous transcripts encoding NUCLEOLIN-like and AGONAUTE-like (AGO) proteins, large ribosomal subunits, and Heat Shock Protein 70 (HSP70) and HSP90; similar genes defined the leaf ridge cluster (DR_C6), along with *ALOG-D1* that regulates spikelet architecture^3^. At LP, the spikelet primordium cluster (LP_C10) was defined by transcripts encoding an Aux/IAA protein (IAA13), SUCROSE SYNTHASE (SUS2), and a peptidyl prolyl cis-trans isomerase, while the spikelet boundary cluster (LP_C6) was defined by genes encoding AGO, HSP70, HSP90, SQUAMOSA-promoter binding protein-like13 (SPL13) and a cytochrome P450 monooxygenase (CYP78-16) that influences organ size^27,28^. The meristem regions were defined by transcripts encoding histones and ribosomal sub-units, and these were maintained between DR and LP. The rachis and basal inflorescence regions of DR were defined by transcripts encoding MT2^29^, the aquaporin TIP1;1^30,31^, a cytoplasmic glyceraldehyde-3-phosphate dehydrogenase (GAPDH) enzyme involved in glycolysis (GapC3)^32^, and homologues of the lectin protein EULS3^33^. Similar transcripts were detected in these regions at LP, as well as transcripts encoding photosynthesis-related proteins (e.g., FERREDOXIN and CHLOROPHYLL A/B BINDING PROTEINS), which is consistent with the onset of rachis greening that occurs during this stage^7,34^.

### Integration of spatial clusters and marker genes highlights domain conservation and differentiation between DR and LP

Next, we investigated the association between clusters within each stage, as well as the relationships between clusters of similar tissue types across stages (Fig. 2a,b). Meristem domains at double ridge were comprised of two pairs of correlated clusters, DR_C2 and DR_C8 and DR_C1 and DR_C10, which localised to apical and lateral regions of the developing inflorescence. Similarly, the well-correlated LP meristem clusters of LP_C3, LP_C4, LP_C9 and LP_C11 localised to the apex of both inflorescence and lateral meristems; together, these data suggest that the position of meristems at apical and lateral regions of the developing inflorescence is conserved between DR and LP. The composition of marker genes and their ontologies within meristem clusters overlapped substantially across stages, indicating their identity is well maintained between DR and LP, and meristems are sustained as cells differentiate into spikelet primordia. The meristem identity of these clusters is supported by their inclusion of genes known to control meristem size, with transcripts encoding FON2-LIKE CLE PROTEIN1/CLAVATA3 (FCP1/CLV3) and CLV1 localising to inflorescence and lateral meristem cells at DR and LP^35–39^. At DR, *FCP1* and *CLV1* transcripts also localised to the leaf ridge cluster. The pair of clusters that defined the vasculature of the rachis and inflorescence base were strongly correlated at both DR and LP, and a large proportion of transcripts that localised to the vascular regions at LP were detected in corresponding clusters at DR. Together, these results suggest that a key event of DR involves the establishment of central vasculature tissue that will eventually become the rachis and stem segments. The two spikelet-associated domains were represented by strongly correlated clusters of DR_C5 and DR_C6 at double ridge, and by LP_C6 and LP_C10 at lemma primordium. Relatively few genes were shared between these clusters within and across stages, indicating that at least two distinct groups of cells are required to coordinate spikelet formation, and the progression of spikelet development alters the transcriptional profile of these domains. Analysis of marker genes between DR and LP indicates transcripts that localised to the spikelet primordia (LP_C10) were expressed in each of the spikelet- and vasculature-associated clusters of DR, while those localising to the spikelet boundary (LP_C6) associate mostly with the leaf ridge and rachis clusters of DR.

**Figure 2.**
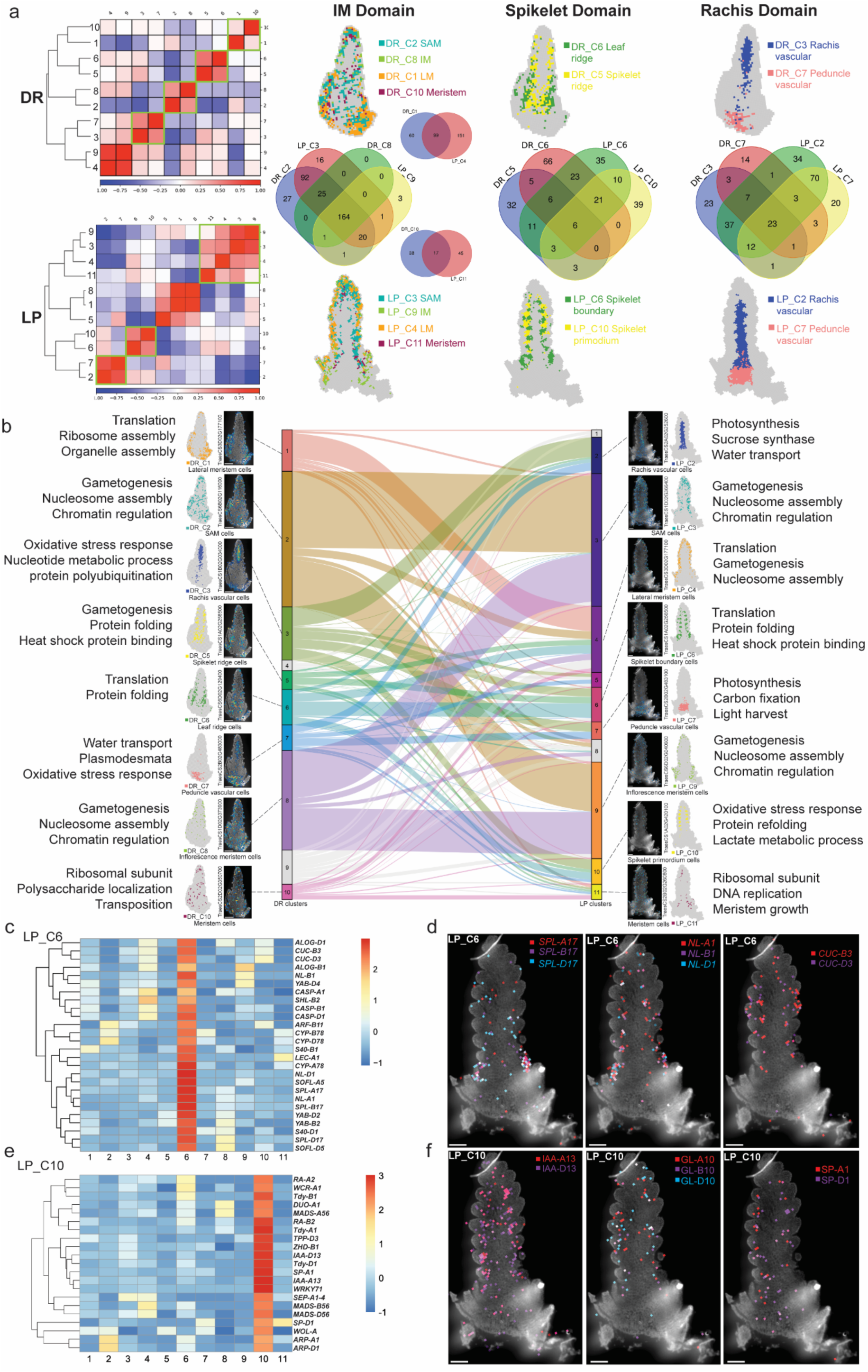
Conserved and divergent spatial domains across developmental stages. **a**, Identification of conserved spatial domains. Pairwise correlation heatmaps and hierarchical clustering define three major cluster bundles: inflorescence meristem (IM), spikelet, and rachis domains. Venn diagrams show the overlap of differentially expressed genes across stage-specific clusters within each domain. Cluster locations are highlighted in representative tissue sections. **b**, Domain continuity and transition between DR and LP. Sankey plot illustrates the continuity and divergence of gene expression programs between DR and LP clusters. Selected marker genes and their enriched Gene Ontology (GO) terms are shown alongside their spatial expression maps. **c,e**, Scaled expression (Z-scores) of genes enriched in LP_C6 (c) and LP_C10 (e). LP_C6 represents a spikelet boundary cluster enriched for *SPL17*, *NL1*, and *CUC3*, while LP_C10 corresponds to spikelet primordia identity with enriched expression of *IAA13*, *GL10*, and *SP1*. **d,f**, Spatial localization of representative genes from LP_C6 (d) and LP_C10 (f), showing distinct expression domains consistent with boundary and spikelet primordia regions, respectively. Scale bars, 200 µm.

To further investigate the identity of the two spikelet-associated domains, we surveyed transcripts expressed in LP_C6 and LP_C10 that were represented by two or three homeologues. All the genes enriched in LP_C6 and LP_C10 and other DR/LP clusters were shown in Supplementary Table 3. Support for the identity of LP_C6 as a spikelet boundary domain came from identification of genes that coordinate lateral organ boundary formation, such as *CUP-SHAPED COTYLEDON3* (*CUC3*)^40^ and *ALOG1* (*ALOG-B1, ALOG-D1*)^3^; ALOG1’s role in boundary formation helps lateral meristems form a single spikelet, rather than a spikelet pair. We also detected transcripts encoding NL1, SPL17 and S40, which suppress bract outgrowth or promote leaf senescence^41–45^. Localisation of these transcripts to the spikelet boundary is consistent with the suppression of leaf primordia that occurs on the abaxial side of developing spikelets, and supports the model proposed in maize whereby SPL17 (ZmTSH4) and NL1 (ZmTSH1) form a network to suppress bract growth and promote lateral meristem indeterminacy^46,47^.

The identification of transcripts encoding a YABBY transcription factor (YABBY6), SHOOTLESS2 (SHL2) and KANADI further links abaxial/adaxial patterning to this cluster. SHL2 controls expression of microRNA 166 (miR166), which targets class III homeodomain-leucine zipper transcription factors, including *HOMEOBOX DOMAIN-2* (*HB-2*), which promotes an adaxial fate and acts antagonistically to abaxial-promoting role of KANADI^12,48–52^. Higher *HB-2* expression promotes formation of a spikelet pair, rather than the typical single spikelet^12^, and transcripts for all three *HB-2* homeologs were enriched in the spikelet primordium (LP_C10). This cluster contained transcripts for all three homeologs of genes encoding a CASPARIAN STRIP MEMBRANE DOMAIN PROTEIN (CASP)^53^, which recruits lignin polymerization machinery for Casparian strip deposition in the endodermis, a SOB FIVE-LIKE (SOFL)^54^ protein involved in cytokinin-mediated development, and CYP78-16^27^ (Fig. 2c,d). This cluster also contained transcripts for three homeologues of *TEOSINTE BRANCHED1 (TB1)*, which regulates spikelet architecture in a dosage dependent manner^11^. The identity of LP_C10 as a spikelet primordium domain is supported by detection of *SEPELLATA1-4* (*SEP1-4*) transcripts, which MERFISH analysis identified as a glume marker^24^. Similarly, this cluster was enriched for genes that regulate spikelet and branch architecture in rice, barley and wheat, such as *MADS56*^55^ and *DUO1*^10^. Three homeologs transcripts were identified for *IAA13*, which controls lateral organ boundary formation in an auxin-dependent manner^56^, and an ACC oxidase (ACO2)^57^ that catalyses the final step of ethylene biosynthesis. The cluster also contained transcripts encoding proteins involved in sugar metabolism and transport, including trehalose-6-phosphate phosphatases (TPP) that localise to comparable regions in maize^58^, and homeologs of *TIE-DYED1* (*Tdy1*)^59^, which encodes a transmembrane protein that loads sugar into phloem (Fig. 2e,f). Together with existing functional evidence, our data suggest that the alternating, distichous arrangement of spikelets along a wheat inflorescence depends on a reticulated lattice-like expression of at least two gene groups, which coordinate either spikelet development or boundary formation.

### Pseudotime analysis reveals developmental trajectories and tissue differentiation in early wheat inflorescence

To investigate tissue differentiation during early inflorescence development, we used Monocle2 to perform pseudotime analysis of cells at DR and LP. Two major developmental trajectories were identified at each stage, with LP displaying a clearer divergence in trajectory topology, consistent with tissue being more differentiated at the later stage. The pseudotime trajectories originated from regions that overlap with meristem clusters and progressed towards those that associate with either spikelets or the rachis (Fig. 3a,c). Trajectory-based clustering of pseudotime-associated cell states further supports the direction of these developmental trajectories. (Fig. 3b,d). DR and LP were each defined by five cell states, with three major states aligning with meristem, spikelet or rachis regions (Fig. 3e-h). Cells from earlier in the trajectory mapped to meristem domains (MD), while more developmentally advanced cells localised to rachis (RD) and spikelet domains (SD). Cells associated with intermediate segments of the trajectories localised to lateral meristem regions, suggesting they contribute to spikelet formation. Next, we performed pseudotime-dependent expression analysis to identify genes that associate with early or late phases of the developmental trajectory. These genes were grouped into four expression profiles, depending on them being active during early (profile 2), late (Profile 1), or at intermediate phases (Profiles 3 and 4) of the pseudotime continuum (Fig. 3i–j; Supplementary Table 4).

**Figure 3.**
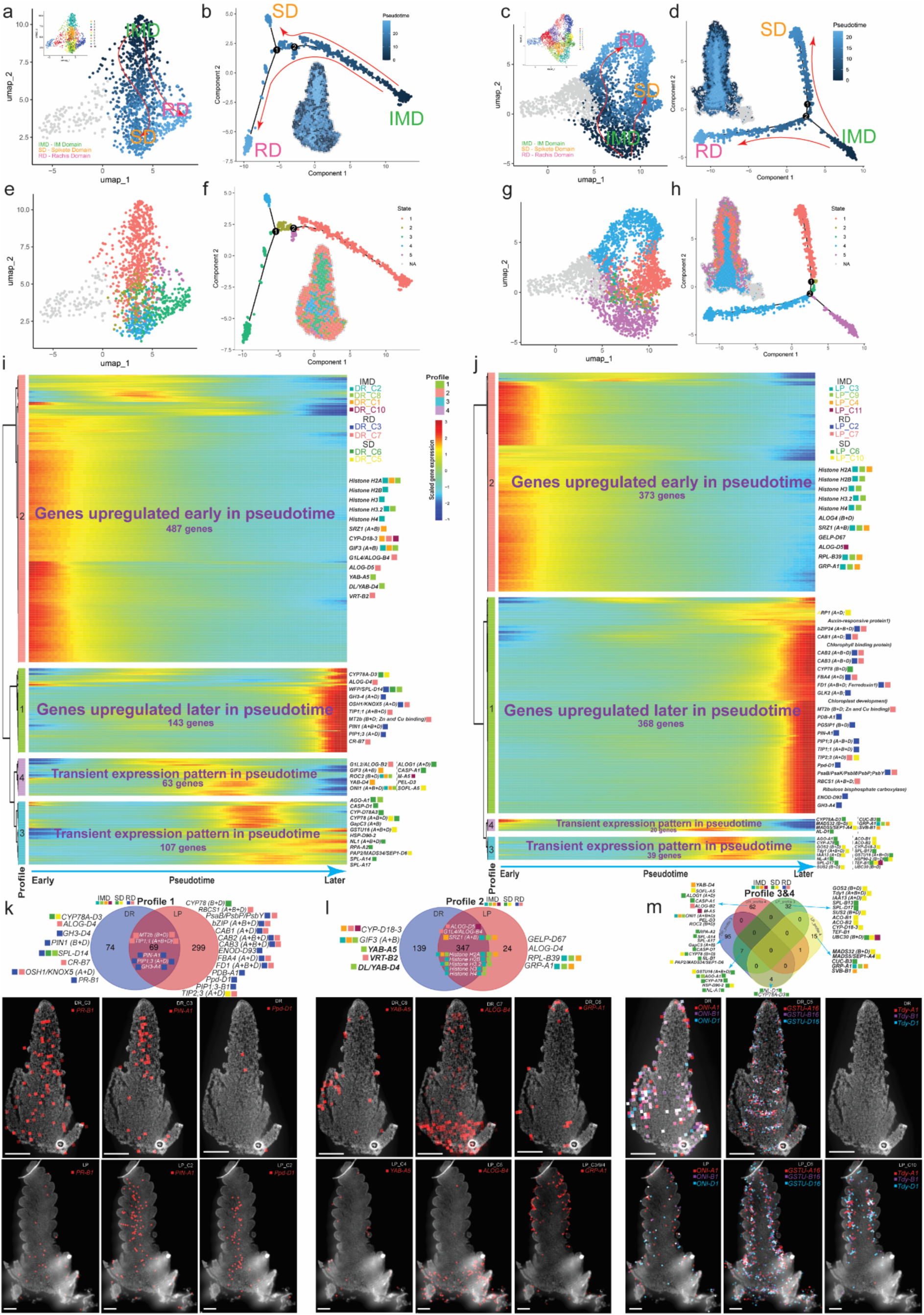
Pseudotime trajectory analysis reveals progressive differentiation of spatial domains in wheat inflorescence. **a–d**, Pseudotime inference using Monocle2 identifies two primary developmental trajectories in DR (a,b) and LP (c,d) stages, originating from the inflorescence meristem domain (IMD) and diverging toward the spikelet domain (SD) and rachis domain (RD). UMAPs are overlaid with pseudotime values (b,d) and domain annotations (a,c), revealing a gradual transition from undifferentiated to differentiated cell populations. **e–h**, Trajectory-based clustering defines seven pseudotime-associated cell states in DR (e) and five in LP (g), reflecting transcriptional heterogeneity. Tissue mapping of these states (f,h) demonstrates spatial coherence of developmental progression. **i–j**, Heatmaps of pseudotime-dependent gene expression profiles in DR (i) and LP (j). Genes were grouped into four profiles based on expression dynamics: Profile 1 – late upregulated; Profile 2 – early upregulated; Profiles 3 and 4 – transiently expressed at intermediate pseudotime. Representative genes are labelled and coloured by their spatial cluster identity. **k–m**, Venn diagrams comparing DR and LP gene sets for each pseudotime profile: late upregulated (k), early upregulated (l), and transiently expressed (m). Examples of shared and stage-specific genes are indicated, including *PR-B1, PIN-A1, Ppd-D1, YAB-A5, ALOG-B4, GRP-A1, ONI1, GSTU16 and Tdy1*. These profiles reveal conserved and distinct regulatory programs controlling phase transitions, spikelet initiation, and meristem identity. Scale bars, 200 µm.

Genes encoding histone and ribosomal proteins are upregulated early in the pseudotime at DR and LP, consistent with meristems featuring early in the developmental trajectory (Profile 2). Many of these genes shared the same profile at DR and LP, supporting the view that meristems are sustained during early inflorescence development (Fig. 3l). Genes encoding MT2b, TIP1;1, GH3-4, PIN1, and the aquaporin PIP1;3, which localise to the rachis, were upregulated late in the DR trajectory (Profile 1) (Fig. 3k; Supplementary Figure 14). A similar gene set is expressed late in the LP trajectory, along with 299 unique transcripts that are enriched in the rachis, indicating more advanced differentiation of vascular tissue at the later stage. The additional genes include *Ppd-D1*^6^*, PHOTOPERIOD-1*-*DEPENDENT bZIP* (*PDB1*)^3^, *basic leucine zipper (bZIP) transcription factor* (*FD1*)^60^ that control flowering time and spikelet number, and those encoding aquaporins (e.g., TIP2;3) and photosynthesis-related proteins (e.g., CAB1/2/3; Psa and Psb proteins) (Fig. 3k; Supplementary Figure 14). Transcripts expressed during transient phases of the trajectory (Profile 3 and 4) of DR and LP are enriched for spikelet development genes (e.g., *ALOG1*, *CYP78*, *IAA13*, *SPL17*, *CUC3*, *MADS32*), with many showing stage- and profile-specific expression, suggesting they perform temporally specific roles during spikelet development (Fig. 3m; Supplementary Figure 14). Together, these data indicate that spikelet development involves dynamic transcriptional changes in cells that differentiate from meristem domains, during stages accompanied by rapid differentiation of central vascular tissue.

### Spatial co-expression network analysis identifies region- and stage-specific gene modules

Next, we performed co-expression network analyses to investigate spatially distinct gene regulatory modules within the inflorescence. Hierarchical clustering of gene expression correlations resolved the LP data into six discrete modules (Fig. 4a; Supplementary Table 5). Modules derived from DR-expressed genes did not separate clearly due to homogenous expression patterns across the inflorescence (Supplementary Figure 13), which is indicative of the undifferentiated tissue at the earlier stage. The six LP modules localised to regions aligned to either central and basal vascular regions (Module 1 (M1), M3 and M5), meristems (M2), or spikelets (M4 and M6) (Fig. 4b,c). The spikelet-associated modules localised to either the central region of the inflorescence, where spikelet formation is most advanced (M4), or to the inflorescence apex where spikelets are less differentiated (M6). Module_4 comprised genes expressed in spikelet primordia and boundary regions, including *TDY1*, *NL1*, and *CYP-D78*, as well as vascular-associated genes such as *PIN1*^61^ and *ENOD93*^62^ (Fig. 4d). This suggests that advanced development of spikelets in the centre involves coordinated regulation of genes across spikelet and vascular tissues. We also detected *NL-A1* and *CYP-D78A3* transcripts at the spikelet boundary during DR, and *ENOD-D93* and *PIN-D1* in vascular tissue at DR, indicating this gene network is involved in early phases of spikelet development (Supplementary Figure 14). Module_6 included spikelet primordia genes such as *MADS32* and the glume marker, *SEP1-4*, as well as genes from meristem clusters (*SVB-1* and *RPL10*), which reflects the ongoing differentiation of spikelets from meristematic tissue in this region (Fig. 4c).

**Figure 4.**
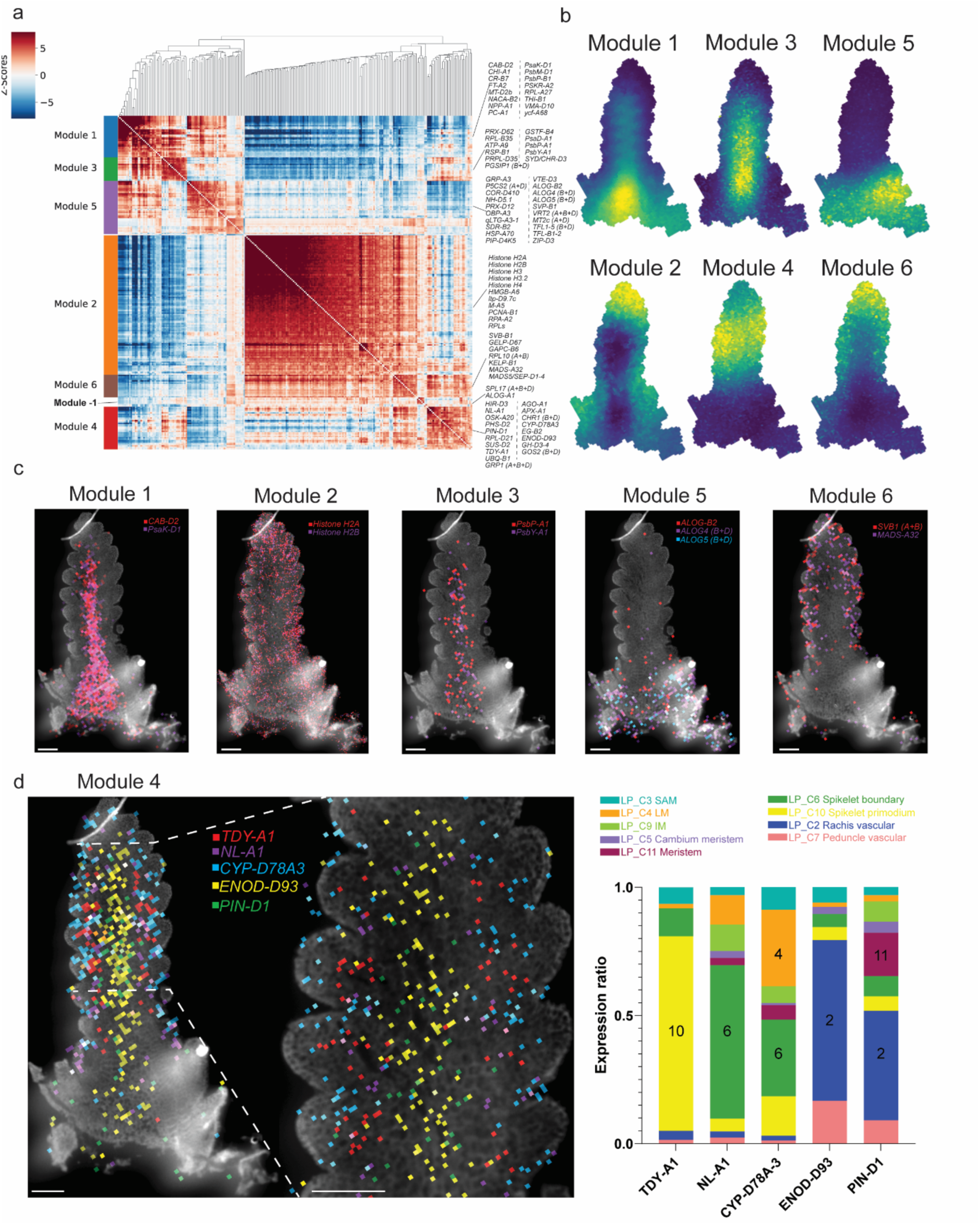
Spatial co-expression modules reveal transcriptional programs that distinguish developmentally diverse regions of the inflorescence. **a**, Co-expression network analysis. Heatmap showing pairwise gene–gene correlations across spatial bins identifies six co-expression modules with distinct expression patterns. Modules enriched for known genes are annotated. **b**, Spatial localization of co-expression modules. Module expression scores are mapped onto DR and LP sections, revealing spatially distinct domains. **c**, Expression of selected module-specific genes shown as overlays on tissue images. **d**, Spatial localization and cluster-specific expression of Module 4 genes. Left, spatial expression of five representative Module 4 genes (*TDY-A1*, *NL-A1*, *CYP-D78A3*, *ENOD-D93*, *PIN-D1*) mapped onto LP sections. These genes are enriched in the central region of the spike, corresponding to the most developmentally advanced spikelet primordia. Right, stacked bar plots show the proportional expression of each gene across LP spatial clusters, with numeric labels indicating the dominant contributing cluster. Module 4 genes are prominently expressed in LP_C10 (spikelet primordium), LP_C6 (spikelet boundary), and LP_C2 (rachis vascular), indicating roles in spikelet differentiation, boundary formation, and vascularization during advanced stages of spikelet development. Scale bars, 200 µm.

Interestingly, the two spikelet modules correlated with a set of spikelet boundary genes (Module_-1), including *ALOG1* and *SPL17* (Fig. 4a), which supports the identification of two discrete regions that contribute spikelet formation along the wheat inflorescence. Module_2 comprised of histone and ribosomal proteins representative of meristem regions, and it localised to apical and lateral regions of the inflorescence (Fig. 4a,c). M1 and 3 localised to vascular regions in the inflorescence that each contain transcripts encoding photosynthesis-related proteins; the separate networks detected for the centre and base of the inflorescence indicate distinct identities for these vascular regions, potentially because the basal region will proceed to form stem internodes and the peduncle (Fig. 4b,c). Genes unique to M1 included *FLOWERING LOCUS T2* (*FT2*) (Fig. 4a; Supplementary Figure 14), which contributes to flowering time and termination of spikelet development^7,63^. The inflorescence base was also represented by M5, which was enriched for transcripts encoding SHORT VEGETATIVE PHASE (SVP1, SVP2/VRT2), TERMINAL FLOWER1 (TFL1) and ALOG (ALOG2, ALOG4, ALOG5) transcription factors (Fig. 4b,c; Supplementary Figure 14). This module aligns with regions of the inflorescence where spikelet development is often delayed, relative to the centre, resulting in infertile spikelets; taken together with known roles for SVP and TFL in repressing early floral transition events, this region may help form a boundary between vegetative and reproductive zones of the developing inflorescence^15–18,20,64^ (Fig. 2c). *ALOG2/4/5* transcripts were also detected in rachis and spikelet regions at DR, suggesting they contribute to boundary formation within the inflorescence during early development^3,65^ (Supplementary Figure 14). Together, these modules provide insights into the progressive phases of spikelet development along the inflorescence, which may explain why spikelet fertility and branching phenotypes are more pronounced in the central region.

### *RA2* expression marks a spikelet domain that controls spikelet formation

The identification of two distinct clusters that align with spikelet initiation and development at DR and LP indicated that at least two groups of cells, with different gene expression profiles, are required to direct the alternating, distichous arrangement of single spikelets along the wheat inflorescence. This model is supported by the role of *ALOG1*, which marks the spikelet boundary cluster at both stages and is required to restrict lateral meristem differentiation to a single spikelet^3,65^. To further explore this model and the role of the spikelet initiation (SI) and primordium (SP) clusters in regulating spikelet architecture, we surveyed genes enriched in the SI and SP clusters of DR and LP, respectively, with a focus on triads of homeologues that are expressed strongly in these domains. One triad includes homeologues of a gene encoding a lateral organ boundary (LOB) domain containing transcription factor that is homologous to RAMOSA2 from maize and Vrs4 from barley^66,67^; we named the gene *RAMOSA2* (*RA2*). *RA2* is expressed exclusively in the spikelet initiation (C5) cluster of DR and predominantly in the spikelet primordium (C10) domain of LP (Fig. 5a,b). We then asked if any of the 256 paired spikelet-producing mutants identified from the Cadenza TILLING population contained mutations in a copy of *RA2*^12^. Paired spikelets are a supernumerary spikelet structure characterised by the formation of a secondary spikelet immediately adjacent to and below the typical primary spikelet, and absence of ALOG1 expression facilitates their development^3,6^. Among class II paired spikelet-producing mutants^12^ that form multiple secondary spikelets was *paired spikelet 3* (*ps3; CAD1591*) (Fig. 5c), which formed secondary spikelets at 45.9 ± 3.2% of rachis nodes (Fig. 5d). The paired spikelets formed predominantly in the centre of the inflorescence, and scanning electron microscopy of developing inflorescences showed the secondary spikelets emerged at the floret primordium and terminal spikelet stages, consistent with our previous analyses (Supplementary Figure 15)^6,11,12^. The *ps3* mutant contained a missense mutation (A91T) in the D genome copy of *RA2*, named *RA-D2* (*TraesCS3D02G093500*) (Fig. 5e); A91 residue is conserved among LOB domain transcription factors, and localises to an alpha helix (α4) that stabilises the homodimerization of RA2 required for DNA binding^68^. The *ps3* line was crossed to *cv.* Cadenza four times to filter background mutations, and BC_3_F_2_ segregating populations were analysed using exome capture sequencing and marker assays to show the *ra-D2* missense allele associated significantly with paired spikelet production (Extended Data Fig. 1). This association was further verified using recombinant inbred lines, which excluded mutations linked to *ra-D2* (A91T) in the exome capture sequence analysis (Extended Data Fig. 1). To support this role for *RA-D2* during spikelet development, we investigated an independent mutant line (*CAD0289*) that contained a missense mutation (L72F) that localises to a neighbouring alpha helix of a GAS (Gly-Ala-Ser) motif that is also required for homodimerization and stability of protein-DNA interactions (Fig. 5e)^68^. Like *ps3,* this *ra-*D2 missense (L72F) allele was only detected in paired spikelet-producing plants of BC_2_F_2_ segregating families, and not in wild type plants that expressed the reference allele (Extended Data Fig. 2).

**Figure 5.**
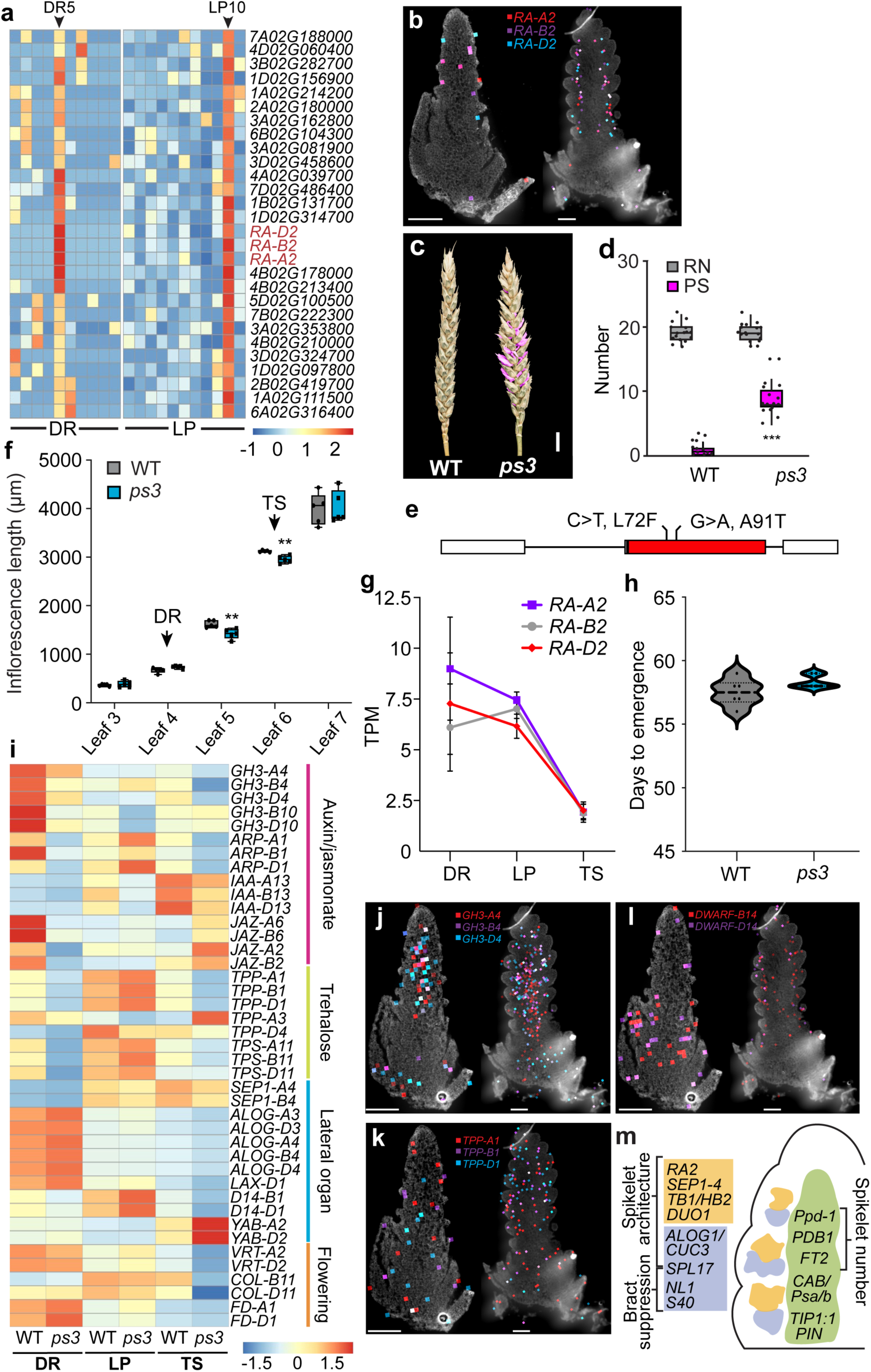
Analysis of developmentally conserved spikelet primordium genes identifies *RAMOSA2* (*RA2*) as a key spikelet regulator. **a**, Heatmap showing genes co-enriched in DR_C5 and LP_C10 clusters that define spikelet primordium domains DR and LP, which included three homeologues of *RA2* (*RA-A2*, *RA-B2*, *RA-D2*). Expression values are scaled by row (Z-score). ‘TraesCS’ is the prefix to each gene ID. **b**, Spatial expression profiles of *RA2* homeologues at DR and LP. Scale bar, 200 μm. **c-d**, Spikes of the *ps3* mutant form multiple secondary spikelets (pink), relative to wild-type (WT) siblings, without altering rachis node numbers (grey). Scale bar, 1 cm. **e**, Schematic of *RA-D2* showing two identified missense mutations (L72F, A91T) that promote paired spikelet development (white, UTR; red, exon 1). **f**, Analysis of inflorescence growth rate and developmental progression in WT (grey) and *ps3* (blue). In the box plot, each box is bound by the lower and upper quartiles, the central bar represents the median, and the whiskers indicate the minimum and maximum values of 4 biological replicates. **g**, Expression of *RA2* homeologues (*RA-A2*, *RA-B2*, *RA-D2*) during early inflorescence development. **h**, Flowering time analysis of *ps3* (blue), relative to WT (grey), shown as a violin plot with individual data points. **i**, Heat map displaying DETs in inflorescences of *ps3*, relative to WT. **j-k**, Localization of selected DETs in DR and LP inflorescences. **m**, A model illustrating how spatially distinct domains regulate spikelet architecture, bract suppression, and spikelet number in bread wheat. In **a** and **i**, normalized expression values are scaled by row (Z-score). DR, double ridge; LP, lemma primordium, TS, terminal spikelet. Asterisks indicate statistical significance (** P< 0.01; *** P< 0.001).

To investigate the effect of these *ra-D2* missense alleles on inflorescence development, we analysed inflorescence growth during early stages when spikelets and florets form. Inflorescence growth was delayed between DR and LP in plants of both mutant lines, relative to wild-type siblings (Fig. 5f). The delay between these stages is consistent with expression of *RA-D2* and its homeologues, *RA-A2* and *RA-B2* (*TraesCS3A02G093200* and *TraesCS3B02G108500*), peaking during DR and LP before declining at TS, with no transcripts detected at later stages (Fig. 5g). The delay was specific to these stages, as mutant plants reached TS and flowered at the same time as wild type, consistent with there being no difference in rachis node number between wild-type and mutant genotypes (Fig. 5f, h; Extended Data Fig. 2).

Next, we performed RNA-seq transcriptome analysis during early developmental stages to investigate molecular processes underlying paired spikelet development in *ps3*. We detected 763 differentially expressed transcripts (DETs) at DR, of which 84.5% were down-regulated in *ps3*, relative to wild type, while 368 and 792 DETs were detected at LP and TS, respectively, with similar proportions of up- and down-regulated genes (Supplementary Table 6). At DR and TS, down-regulated transcripts were enriched for processes related to trehalose metabolism, consistent with transcriptome changes detected in the barley *vrs4* mutant (Fig. 5i; Supplementary Figure 16)^67^. Downregulated transcripts included homeologues for three genes encoding trehalose 6-phosphate phosphatases (TPP; TPP1, TPP3 and TPP4), and an alpha-trehalose-phosphate synthase (TPS11). Other downregulated transcripts in *ps3* at DR include those encoding a SEPALLATA-like MADS box transcription factor (SEP1-B4), and proteins involved in auxin and jasmonate signalling processes, such as Gretchen Hagen3 (GH3) acyl acid amido synthetases (e.g., GH3-8) and jasmonate ZIM-domain proteins (e.g., JAZ2, JAZ6). Enrichment of genes involved in auxin signalling pathways was also detected among downregulated transcripts at LP and TS, including those encoding auxin/indole-3-acetic acid proteins (AUX/IAA; IAA13, IAA26), GH3-1, GH3-4, an auxin repressed protein (e.g., ARP1) and a small auxin up-regulated RNAs (SAUR). Transcripts downregulated in *ps3* at TS were also enriched for processes related to lateral organ formation and strigolactone signalling, such as *ALOG3, ALOG4, LAX1, WHEAT ORTHOLOGUE OF APO1* (*WAPO1*), and *DWARF14* (*D14*)^25,65,69–72^. Similarly, genes involved in flowering and spikelet development (e.g., *VEGETATIVE TO REPRODUCTIVE TRANSITION2* (*VRT2*)^15–18,20^, *FLOWERING LOCUS D1* (*FD1*)^60^, *CONSTANS-like 10* (*COL10*)^73^) were significantly downregulated in *ps3* at TS, relative to wild-type, while YABBY2 transcripts were significantly higher in *ps3*. Curiously, no transcripts that are expressed in spikelet boundaries and proposed to act antagonistically to *RA2* were significantly mis-regulated in *ps3,* relative to wild-type; nonetheless, all homeologs for *SPL17, NL1,* and *LG2*, *CUC3* and *ALOG1* were moderately higher in *ps3* at DR, relative to WT, and lower at TS, indicating that delayed differentiation of the spikelet meristem did influence genes expressed in the neighbouring boundary region (Fig. 5i; Supplementary Table 7). Together, these results indicate that secondary spikelet formation in *ps3* involves mis-regulation of processes involved in trehalose and hormone metabolism and signalling, as well as flowering and lateral organ formation.

To further investigate the DETs, we used the spatial transcriptome data to localise their expression within the developing inflorescence at DR and LP. Among the DETs detected in *ps3* at DR, 24.7% (107/434) of genes localised to the spikelet-associated clusters, while 26.7% (116/434) and 25.1% (109/434) were from the meristem and rachis clusters, respectively. At LP, a similar proportion of DETs localised to the spikelet (21.7%; 54/249) and rachis (26.1%; 65/249) associated clusters, while fewer were detected in meristem regions (14.1%; 35/249). Most of the differentially expressed *TPPs* (6/7 genes) and *TPS-B11* localised to the spikelet primordium cluster at LP, while there was no shared location for these transcripts at DR. DETs encoding proteins involved in auxin signalling and metabolism, such as GH3-4, ARP1, IAA13, also located to the spikelet primordium clusters, as did LAX1. Transcripts encoding the YABBY transcriptions factors (e.g., YAB2), LAX1, SPL17 and D14, were divided between the spikelet primordium and boundary cells. Transcripts encoding proteins involved in flowering (COL10) and auxin transport (e.g., PIN10) were present in rachis and peduncle tissue, while *VRT2*, *ALOG3*, *ALOG4* and *JAZ6* transcripts localised to the inflorescence base. Taken together with the bulk tissue RNA-seq analysis, the spatial transcriptome reveals that many transcripts differentially expressed in *ps3* co-localize with *RA2* expression in spikelet-associated clusters, which is consistent with the modified spikelet architecture observed in this genotype.

## DISCUSSION

Our stereo-seq transcriptome analysis provides a high-resolution, state-of-the-art spatiotemporal map of gene expression during spikelet initiation and differentiation in the wheat inflorescence. By localizing transcripts within intact inflorescence sections at double ridge and lemma primordium, we identified distinct gene expression domains that correspond to developing spikelets, apical and lateral meristems, and the central vascular regions of the rachis and inflorescence base. Together with functional knowledge of genes that localise to these domains^6,10–12,15–18,20,42,72^, the spatial transcriptome map provides new insights into how gene expression is coordinated across different cell types to determine the architecture, number and fertility of spikelets that form along the wheat inflorescence (Fig. 5m).

In regions that aligned with developing spikelets, we identified two distinct clusters at DR and LP that were arranged in an alternating distichous pattern flanking the central rachis. One these clusters specifies a group of cells that form spikelet primordia, characterised by expression of *RA2*, *SEP1-B4*, *IAA13* and *DUO1*, while the other marks a boundary region, enriched for transcripts of *SPL14/17*, *NL1*, *S40,* and *ALOG1* that perform roles in bract suppression and spikelet development^41–47,72^. We propose that these two groups of cells function synergistically to coordinate the formation of short lateral branches, each composed of a single spikelet that lacks a subtending bract, and their reiterative expression pattern along the inflorescence facilitates the sequential formation of multiple rachis nodes (Fig. 5m). In this model, spikelet boundary cells perform two distinct roles in wheat. First, they are likely to suppress bract outgrowth through the activity of *NL1*, consistent with findings from corresponding mutants in barley, rice, and maize^43,46,74,75^. Second, they promote formation of a single spikelet—rather than a spikelet pair—in the adjacent primordium, supported by evidence that *ALOG1* functions non-cell-autonomously to suppress paired spikelet development in wheat and barley^65,72^. The proposed role that boundary cells perform in generating multiple rachis nodes through there reiterative expression pattern along the inflorescence is supported by *spl14* and *spl17* wheat mutants forming significantly fewer spikelets than wild-type siblings, with many basal spikelets aborting their development^42^. The localisation of three *RA2* homeologs to the spikelet primordium cluster at DR and LP indicates it is a key gene driving spikelet initiation and development. This role is supported by *ra-D2* mutants forming paired spikelets, which form when differentiation of a lateral meristem into a spikelet is delayed^6,11^–the role of RA2 in spikelet differentiation is also supported by studies in maize and barley^66,67^. Taken together with the functional characterisation of other paired spikelet mutants, we propose that genes expressed in the spikelet primordium and boundary regions act cooperatively to determine spikelet architecture, facilitating formation of single spikelets on opposite sides of the central rachis in an alternating phyllotaxy (Fig. 5m)^6,10–12,72^.A defining feature of the wheat inflorescence is that central spikelets produce more fertile florets than those at the apex or base, which is associated with spikelets developing earlier in the centre than the top or bottom^19,76–78^. The spatial transcriptome data and gene co-expression network analysis provided valuable insights into the molecular basis of asynchronous spikelet development, with separate modules identified for the centre, apex and base. The two modules representing the centre and apex included genes from spikelet and rachis tissues; however, the apical module was enriched for transcripts corresponding to an earlier phase of the pseudotime trajectory compared to than those observed in the central module. These results suggest that apical spikelets share a similar transcriptome profile to central spikelets, but are developmentally delayed. The inflorescence base was specified by a module that included genes such as *VRT2*, *SVP1* and *TFL1*, which restrict the vegetative to reproductive transition and associate with rudimentary spikelet formation at the inflorescence base^15–18,20^. These results suggest that central spikelets are more fertile because they are more developmentally advanced than apical spikelets at the lemma primordium stage, while the delayed development of basal spikelets appears to result from expression of *SVP* and *TFL* genes in adjacent cells.

In addition to genes expressed in spikelet-associated regions, transcripts for other known regulators of inflorescence architecture localised to the central vasculature of the rachis and the inflorescence base. Curiously, these regulators–including *Ppd-1*, *FT2,* and *PDB1*–are known to control spikelet number rather than their configuration, indicating that genes expressed in the vasculature affect rachis node number (Fig. 5m) ^6,7,63,72,79,80^. This model is supported by the enrichment of transcripts encoding photosynthesis-related proteins in the rachis vasculature, as reduced chlorophyll accumulation caused by absence of the vascular-expressed *HvCMF4* (*CCT MOTIF FAMILY4*) restricted final spikelet number and grain production in barley^81^. In support of this model, we observed that transcripts encoding photosynthesis-related proteins accumulated in the rachis during LP, which is when rachis greening initiates and genetic differences in spikelet number are determined (e.g., *Ppd-1* near-isogenic lines^7^). Interestingly, rachis-associated cells also showed strong expression of transporter-encoding transcripts, including aquaporins and *PIN1*, at both the DR and LP stages. This observation suggests that assimilate and hormone transport plays a crucial role in early inflorescence development and that these transporter genes could be promising targets for increasing spikelet number or fertility in wheat.

A third major gene expression domain identified in our data corresponds to meristematic cells, characterised by the expression of histones, ribosomes, and known meristem regulators such as *FCP1* and *CLV1*^35–39^. Meristem-associated clusters were detected in the outer layers of both apical and lateral regions of the developing inflorescence, consistent with the localised expression patterns of key meristem regulators in maize and barley. These results indicate that similar cell types contribute to spikelet development along its entire length, rather than originating from a single pool of cells at the apex. These clusters consistently expressed histone- and ribosome-encoding genes at both DR and LP stages, suggesting that meristematic cells are maintained during spikelet differentiation, potentially to support the ongoing development of florets and floral organs. The substantial presence of these clusters at LP indicates that depletion of meristematic cells is not the cause of spikelet development terminating; nonetheless, based on evidence from barley and maize, spikelet or floret number could be boosted in wheat by increasing the activity of meristem maintenance genes^37,39^.

In summary, these data provide a valuable resource for identifying genes that regulate inflorescence development in wheat, which would be difficult to obtain using traditional genetic approaches, given the complexity and redundancy of the hexaploid genome. Together with functional gene characterisation, our findings suggest that genes expressed in the spikelet primordium and boundary regions act cooperatively to shape spikelet architecture, while transcripts enriched in the rachis and inflorescence base influence spikelet number and fertility. Given genetic variation underlying yield-related traits has been studied extensively in wheat, we propose that the integration of our spatial transcriptome data with identified quantitative trait loci and genome sequence information will offer a powerful approach to identify candidate genes associated with grain production^6,11,82–85^. In this context, a key advantage of hexaploid wheat is that the expression of all three homeologs in a specific region can serve as a strong filter for identifying candidate genes–an approach supported by our identification of *RA2*. Together, these data provide a crucial framework for deciphering genes and biological processes that govern spikelet number and arrangement in wheat inflorescences. When combined with similar studies in maize, barley and rice, this work may help uncover the genetic basis of inflorescence architecture diversity among our major cereals^22,24,39^.

## Methods

### Plant materials and growth conditions

The bread wheat (*T. aestivum*) cultivar Mace was used for the spatial transcriptome analysis. The *paired spikelet3* (*ps3*) and *CAD0289* mutant lines used in this study were identified from a screen of the hexaploid wheat ethyl methanesulfonate-induced TILLING population (cv. Cadenza), as described previously. The *ps3* near-isogenic lines (NIL) used were derived from *CAD1591*, crossed to *cv.* Cadenza to generate BC_3_F_4-5_ lines that segregated either for the wild-type or mutant allele of *RA-D2*; the ‘wild-type’ genotype was the NIL derived from segregating populations that did not form paired spikelets. A similar approach was used for *CAD0289*, except phenotype analysis was performed using BC_1_F_2-3_ generation. For both *ps3* and *CAD0289*, we analyzed two independent families generated from separate crosses to *cv.* Cadenza.

Plants were grown in glasshouses under long-day (16-hour light/8-hour dark) photoperiods with Heliospectra light-emitting diode lights (Heliospectra, Sweden), with day and night temperatures of 20° and 15°C, respectively. Extralong daylength conditions (to 22-hour light/2-hour dark) were used for the accelerated generation of backcrossed germplasm for *CAD1591* and *CAD0289*.

### Stereo-seq capturing and library construction

The library preparation and sequencing process for Stereo-seq was based on the modified version of the Stereo-seq standard protocol (v.1.1). Embedded tissues were sectioned longitudinally at a thickness of 12 μm using a Leica CM1800 cryostat. The tissue sections were adhered to the surface of the Stereo-seq chip and incubated at 37 °C for 2 minutes. They were then fixed in methanol and incubated at −20 °C for 30 minutes. Subsequently, the sections were stained with Qubit ssDNA stain to label nuclei and Fluorescent Brightener 28 (Sigma, F3543-5g) for cell wall visualization (Supplementary Figure 17). Tissue integrity and sectioning quality were evaluated by image-based quality control under fluorescence microscopy prior to proceeding. The sections were then de-crosslinked in TE buffer (10 mM Tris, 1 mM EDTA, pH 8.0) at 55 °C for 1 hour. Afterward, the sections were permeabilized at 37 °C for 12 minutes to enable efficient mRNA capture by spatially barcoded probes, followed by on-slide cDNA synthesis and incubated overnight at 42 °C for reverse transcription and cDNA synthesis. The tissue was subsequently digested at 37 °C for 30 minutes and treated with Exonuclease I (NEB, M0293L) for 1 hour to remove free nucleic acids. Resulting cDNA libraries were quantified using Qubit and assessed for size distribution using TapeStation. Finally, Pooled libraries were sequenced on the MGI DNBSEQ-G400 platform (with either 50 bp or 100 bp paired-end sequencing), generating a total of 1.76 billion high-confidence reads.

### Stereo-seq data processing and quality control

Stereo-seq raw data processing and quality control were performed using the SAW software suite (available at GitHub - STOmics/SAW). Read 1 contained the coordinate identity (CID) and molecular identifier (MID) sequences, with CID spanning nucleotides 1–25 and MID spanning nucleotides 26–35. Read 2 comprised the cDNA sequence. During the initial processing, CID sequences were mapped to the predefined coordinates of the *in situ* captured chip, allowing for a single-base mismatch to account for sequencing and PCR errors. Reads with MID sequences containing ambiguous ‘N’ bases or more than two bases with quality scores below 10 were excluded. The CID and MID identifiers were appended to the respective read headers.

The retained reads were then aligned to reference genomes (IWGSC RefSeq v1.1) using the STAR aligner. Only mapped reads with a MAPQ score greater than 10 were annotated to their corresponding genes. Unique molecular identifiers (UMIs) with identical CIDs and gene loci were merged, allowing for a single-base mismatch to correct sequencing and PCR errors. The single-stranded DNA photo feature in the SAW workflow enabled accurate mapping of raw reads to their precise positions on the tissue. Finally, an expression count matrix was generated, documenting the expression levels of each gene in every spot.

### Unsupervised clustering analysis of Stereo-seq data

We performed unsupervised clustering analysis using the Seurat R package (v4.4.3). First, we normalized the UMI count matrices using the SCTransform function. Feature genes were then selected with the ‘FindVariableGenes’ function, employing the vst method and retaining 2,000 variable features. Subsequently, we conducted principal component analysis (PCA) using the RunPCA function on the selected variable genes, retaining 30 principal components for dimensionality reduction. Next, we constructed the SNN graph using the FindNeighbors function and performed clustering with the Louvain algorithm via the FindClusters function. The clustering resolution parameters were adjusted according to the specific datasets. For visualization, we used nonlinear dimensionality reduction algorithms (RunUMAP). Batch effects between samples were corrected using the ‘RunHarmony’ function to ensure that observed differences in gene expression are due to biological rather than technical variations.

### Identification of Cluster-Enriched Genes

To identify cluster-enriched genes within the spatial transcriptome data, we used the FindAllMarkers function in Seurat. We applied a two-sided Wilcoxon rank sum test to compare gene expression levels between clusters, setting a log fold change threshold of 1.5 (logfc.threshold = 0.58) to highlight genes with at least a 1.5-fold expression difference. We also required that genes be expressed in at least 25% of the cells within a given cluster (min.pct = 0.25) to focus on biologically relevant markers. Finally, we controlled for multiple comparisons using the Bonferroni correction, retaining only genes with an adjusted P value below 0.05.

### Monocle 2 analysis

To investigate the developmental trajectory of cell clusters in the DR and LP tissues, we employed Monocle2^86^. Specifically, we extracted data from the key clusters: DR_C1, C2, C3, C5, C6, C7, C8, C10 and LP_C2, C3, C4, C5, C6, C7, C9, C10, C11. To identify the most informative genes for this analysis, we utilized the ‘FindAllMarkers’ function from Seurat (v4.4.3) to calculate the top marker genes for each cluster. These marker genes were then selected for subsequent analysis with Monocle2. Dimensionality reduction was performed using the DDRTree method, which helped to reconstruct the developmental trajectory. The resulting trajectory was visualized using the plot_cell_trajectory function. In this analysis, the cluster corresponding to the SAM (DR_C2 and LP_C3) was designated as the starting point. Additionally, we identified a branch point to analyze divergent differentiation pathways. To detect genes that are differentially expressed in a pseudotime-dependent or branch-dependent manner, we applied the BEAM algorithm. Genes that exhibited significant branch dependency were visualized using the plot_genes_branched_heatmap function. This approach allowed us to elucidate the dynamic gene expression changes underlying the developmental progression of the early wheat inflorescence.

### Spatial co-expression module analysis

To identify gene co-expression modules in spatially resolved transcriptomic data, the Hotspot algorithm (v1.1.1) was employed. The spatial transcriptomic data were first converted into a Hotspot object using the Hotspot function. A K-Nearest Neighbors (KNN) graph was then constructed using the create_knn_graph function with n_neighbors set to 300 to capture the spatial relationships between cells. Genes were grouped into co-expression modules using the ‘create_modules’ function, with a minimum gene threshold of 15 and an FDR threshold of 0.05 to ensure the detection of biologically meaningful and statistically significant modules. Finally, ‘calculate_module_scores’ function was used to compute module scores for each cell, identifying cells that highly express the genes within specific modules.

### GO enrichment analysis

To further elucidate the biological functions of the cluster-enriched genes, we performed Gene Ontology (GO) enrichment analysis using the clusterProfiler package (v4.12.6). This analysis identifies overrepresented biological processes, molecular functions, and cellular components among the cluster-enriched genes, helping to uncover the underlying biological themes associated with each cluster. The results of the GO enrichment analysis were visualized using the ggplot2 package, a versatile and widely used R package. Furthermore, to explore the shared cluster specific differential expressed genes between two key developmental stages (e.g., DR and LP), we utilized the networkD3 package to create Sankey plots. It provides a clear representation of the shared differential expressed genes between stages, highlighting potential commonalities and differences in gene expression patterns.

### RNA extractions and RNA-seq transcriptome analysis

RNA was extracted from whole inflorescences at the double ridge (DR), lemma primordium (LP) and terminal spikelet (TS) stages of early inflorescence development. Each sample consisted of a pooled collection of 6-12 inflorescences, stored in DNA/RNA Shield solution (Zymo Research, R1100-50) at −20°C. Three biological replicates were collected for *ps3* and the wild-type sibling line, and cv. Cadenza. Total RNA was extracted using TRIzol/Chloroform (TRI Reagent, Sigma-Aldrich; Chloroform, ≥99.8%, Thermo Fisher Scientific, USA) and purified using the RNA Clean and Concentrator kit (Zymo Research, R1013) following the protocol outlined by Millán Blánquez^87^. RNA concentration was measured using the Invitrogen™ Qubit™ 3 Fluorometer and Qubit™ RNA HS (High Sensitivity) Assay Kit (Invitrogen, Thermo Fisher Scientific, USA), and RNA integrity was examined using the Agilent 5400 Fragment Analyzer (Agilent Technologies, USA). Library construction and RNA sequencing were performed by Novogene (Novogene HK Company Ltd., Hong Kong). Sequencing libraries were generated using the NEBNext Ultra RNA Library Prep Kit for Illumina (NEB), and index codes were added to attribute sequences to each sample. For sequencing, clustering of the index-coded samples was performed on the cBot Cluster Generation System using the TruSeq PE Cluster Kit v3-cBot-HS (Illumina, USA). After cluster generation, the libraries were sequenced on an Illumina NovaSeq platform to generate 150-bp paired-end reads.

Raw reads were trimmed and filtered using fastp tool^88^ (v0.20.0), and cleaned reads were pseudo-aligned to the IWGSC Chinese Spring gene model index v1.1 using kallisto^89^ (v 0.46.1), as described by Ramírez-González *et al.* (2018)^90^. On average, over 77% of raw reads were successfully pseudo-aligned to the reference transcriptome and used for transcript quantification. Differential gene expression analysis was performed using the R package DESeq2^91^ (v1.49.3) with default parameters. Only transcripts/genes with expression level ≥ 0.5 transcripts per million (TPM) in one sample at least were included. Transcripts/genes with a false discovery rate (FDR) adjusted *P* value (*q* value) < 0.05 were considered differentially expressed. These were further categorized into lists of up- and down-regulated genes at the DR, LP, and TS stages in *ps3* for each pairwise comparison, relative to the wild-type sibling, based on *q* value < 0.05 and mean TPM fold change > 0.5.

### Fine mapping of the *ra-D2* allele in *ps3* mutant lines

To fine map the causal mutation in *ps3*, two independent segregating BC_2_F_2_ families were generated from a cross between *CAD1591* and *cv.* Cadenza. Two bulk DNA samples were generated for each of the two families: the wild-type bulk included DNA of individuals that produced normal inflorescences without paired spikelets, and the mutant bulk included DNA of individuals that produced paired spikelets (> 8 secondary spikelets per inflorescence). Genomic DNA was extracted from leaves, as described previously^12^. Exome capture sequence analysis was performed as described previously^12^, with identified SNPs compared between the two families to eliminate non-causal variant alleles and SNPs were verified by cross-referencing the identified alleles to those identified previously for *CAD1591*^92^. Loci on chromosomes 3D and 6A were identified as potential regions for the causal mutation.

To refine the mutation site, paired spikelet-producing BC_2_F_3_ plants were crossed to Cadenza, and self-fertilized to generate BC_3_F_2_ families. Individual lines were phenotyped for paired spikelets and genotyped using Kompetitive allele–specific PCR (KASP)–based markers for the SNPs identified using exome capture analysis. This analysis identified a region between chr3D.39387558 – 76931611 as the unique locus that associated with paired spikelet production. Individuals that displayed heterozygosity within the region on chromosome 3D were used to generate homozygous recombinant inbred lines (RILs) with different segments of the identified locus. Subsequent phenotype analysis of lines containing either variant or reference alleles for each site across the locus defined marker chr3D.47494317 (G > A; *TraesCS3D02G093500*) as a unique allele that associated with paired spikelet production in all lines. The sequence of *TraesCS3D02G093500* was examined in wild-type and mutant NILs using genomic DNA and complementary DNA, which confirmed a G > A mutation at 271 bp of the coding region for *TraesCS3D02G093500*.

### Phenotypic analyses

Rachis nodes (primary spikelets) and secondary spikelets were recorded for inflorescences of the main stem and first tiller, as described previously^6^. Secondary spikelet distribution was determined as described previously^6^, using inflorescences from the main stem. The rate of inflorescence development was determined for the *ps3* and CAD0289 mutant lines, relative to their respective wild-type siblings, by measuring inflorescence length at intervals defined by leaf emergence (leaf 4 to 7). Flowering time was determined for each genotype at the emergence of the inflorescence from the boot. Scanning electron microscopy was performed on developing inflorescences of wild-type and *ps3* mutants at the late terminal spikelet stage, as described previously^11^.

## Acknowledgement

We thank Dr. Xiujuan Yang (The University of Adelaide) for her advice and guidance on sample preparation for spatial transcriptomics, and we acknowledge Dr. Nick Li (Decode Science; formerly a Field Application Scientist at BGI Research) for his technical support related to the Stereo-seq platform. This work was supported by funding from the Grains Research and Development Corporation, the Australian Research Council (FT210100810), the Royal Society (UF150081), the Biotechnology and Biological Sciences Research Council (BB/T007133/1), and Biological Breeding-National Science and Technology Major Project (2023ZD04073).

## Author contributions

Y.Q. and C.T. contributed equally to this work. Y.Q. and S.B. conceived the study and wrote the manuscript. Y.Q. and M.I. performed sample preparation and conducted the Stereo-seq experiments. Y.Q. and C.T. carried out the spatial transcriptomic analyses with the help of L.Y. and J.S. S.B. supervised the project and led the *RA2* functional study with support from Y.Q., M.P., A.K.A., and S.S. All authors contributed to the review and editing of the manuscript and approved the final version for submission.

## Extended data

**Extended data Fig. 1.**
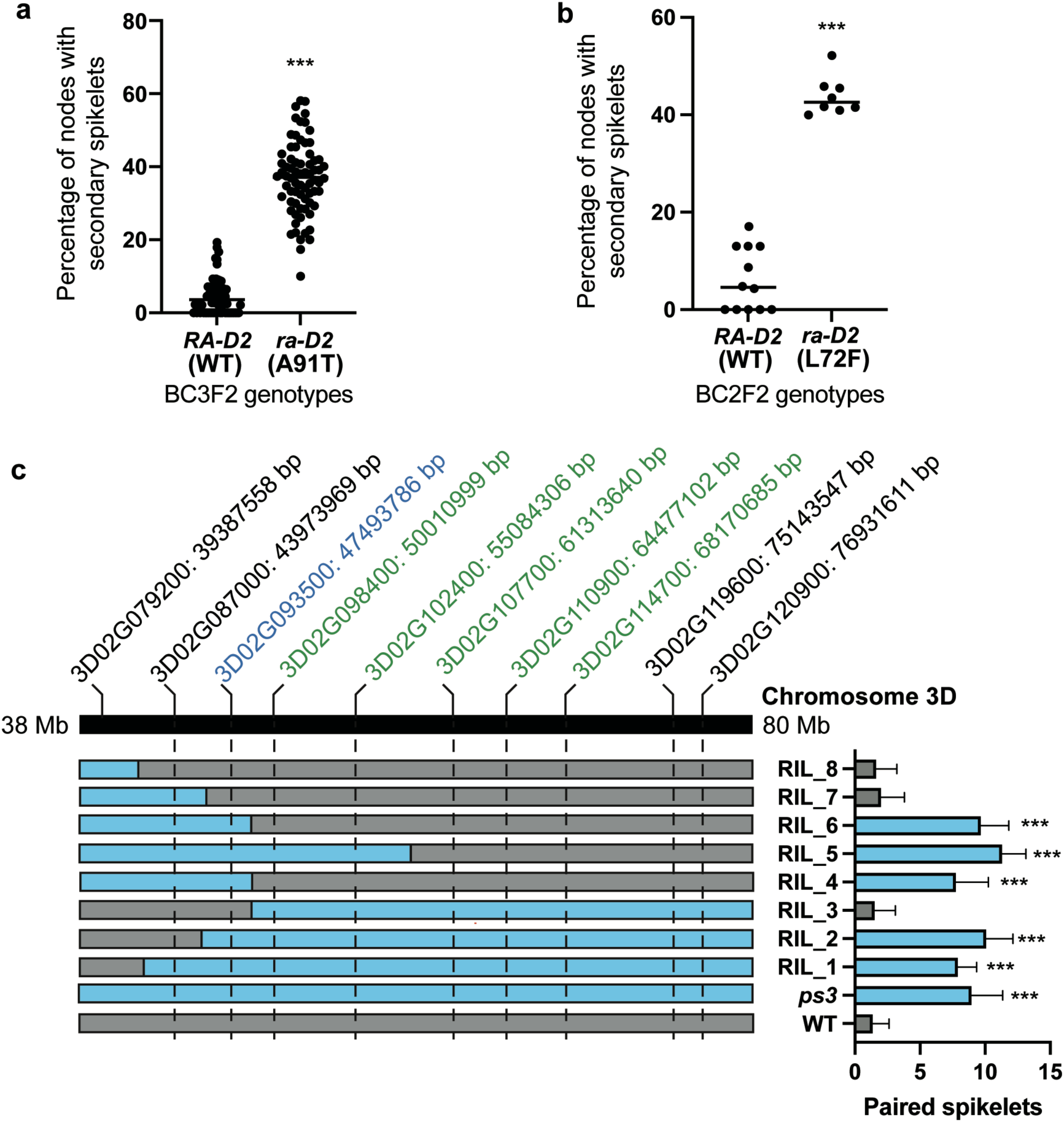
Mapping and validation of the *ps3* locus associated with paired spikelets. **a-b**, quantification of paired spikelet formation in segregating BC3F2 and BC2F2 populations carrying either WT or mutant alleles of *RA-D2* (A91T and L72F, respectively). Each dot represents a biological replicate. ***p < 0.001 (two-sided t-test). **c**, fine-mapping of *ps3* using recombinant inbred lines (RILs). Genotypes are shown across Chromosome 3D from 38 to 80 Mb using KASP markers (listed above, with physical positions). Blue bars indicate *ps3* mutant-derived segments. The number of paired spikelets in each line is shown to the right. Lines carrying the mutant allele exhibit significantly increased paired spikelets (***p < 0.001).

**Extended data Fig. 2.**
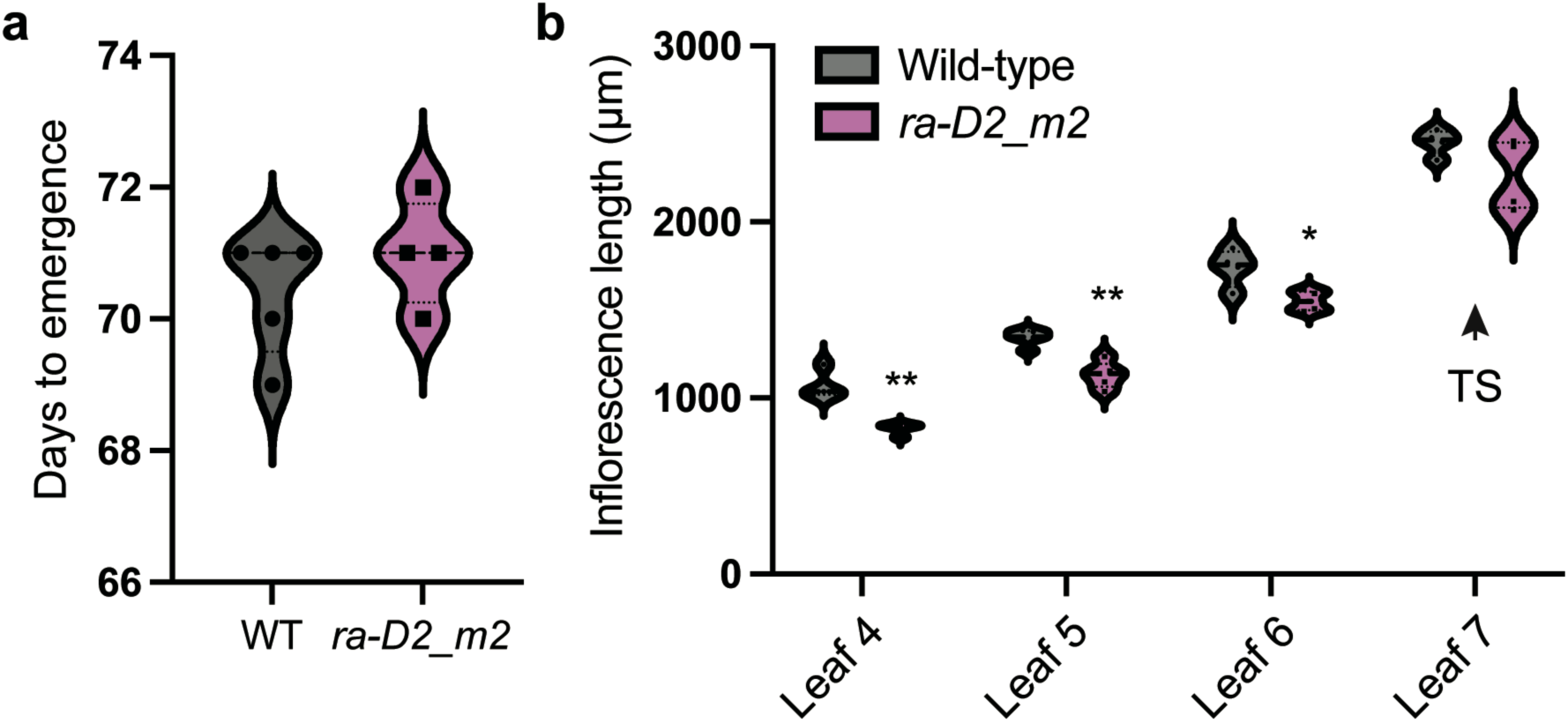
Developmental analysis of *CAD0289* (*ra-D2_m2*) mutant. **a**, days to emergence are not significantly altered in *ra-D2_m2* compared to WT. **b**, Inflorescence length measured at successive leaf stages (leaf 4 to leaf 7) reveals reduced growth in *ra-D2_m2* during early stages, with partial recovery at terminal spikelet (TS) stage. Asterisks indicate statistical significance (*p < 0.05, **p < 0.01; two-sided t-test).

## Supplementary materials

**Supplementary Figure 1. Overview of spatial resolution at different bin sizes in double ridge (DR) stage sections.** Spatial maps and UMAP projections of DR stage sections showing cluster resolution at bin sizes including Bin20, Bin40, Bin50, Bin60, and Bin80.

**Supplementary Figure 2. Overview of spatial resolution at different bin sizes in lemma primordia (LP) stage sections.** Spatial maps and UMAP projections of LP stage sections showing cluster resolution at bin sizes including Bin20, Bin40, Bin50, Bin60, and Bin80.

**Supplementary Figure 3. Validation of DR_C1 to C3 marker genes with MERFISH data.** Comparison of gene expression profiles for DR cluster 1 to 3 between spatial transcriptomics and publicly available MERFISH data, showing consistent spatial localization. Genes including *YAB4*, *TraesCS4D02G022500*, *TraesCS4D02G076900*, *KNOX5*, and *MT2b*. Scale bars, 200 µm.

**Supplementary Figure 4. Validation of DR_C4 to C6 marker genes with MERFISH data.** Comparison of gene expression profiles for DR cluster 4 to 6 between spatial transcriptomics and publicly available MERFISH data, showing consistent spatial localization. Genes including *GRF3*, *CUC3*, *OMTN5*, *ROC4*, and *AG2*. Scale bars, 200 µm.

**Supplementary Figure 5. Validation of DR_C6 to C8 marker genes with MERFISH data.** Comparison of gene expression profiles for DR cluster 6 to 8 between spatial transcriptomics and publicly available MERFISH data, showing consistent spatial localization. Genes including *NL1*, *ALOG4*, *ALOG5*, *MT2b*, and *LSY1*. Scale bars, 200 µm.

**Supplementary Figure 6. Validation of DR_C8 to C10 marker genes with MERFISH data.** Comparison of gene expression profiles for DR cluster 8 to 10 between spatial transcriptomics and publicly available MERFISH data, showing consistent spatial localization. Genes including *MADS32*, *ROC4*, *SCR2*, *ONI1*, and *TraesCS7A02G341800*. Scale bars, 200 µm.

**Supplementary Figure 7. Validation of LP_C1 to C3 marker genes with MERFISH data.** Comparison of gene expression profiles for LP cluster 1 to 3 between spatial transcriptomics and publicly available MERFISH data, showing consistent spatial localization. Genes including *ROC3*, *Ppd1*, *CAB3*, *SEP1-5*, and *AGO14*. Scale bars, 200 µm.

**Supplementary Figure 8. Validation of LP_C4 and C5 marker genes with MERFISH data.** Comparison of gene expression profiles for LP cluster 4 and 5 between spatial transcriptomics and publicly available MERFISH data, showing consistent spatial localization. Genes including *CUC1*, *ROC7*, *YAB4*, *MND1*, and *VRT2*. Scale bars, 200 µm.

**Supplementary Figure 9. Validation of LP_C6 and C7 marker genes with MERFISH data.** Comparison of gene expression profiles for LP cluster 6 and 7 between spatial transcriptomics and publicly available MERFISH data, showing consistent spatial localization. Genes including *GRF9*, *NL1*, *SPL17*, *YAB6*, and *CAB1*. Scale bars, 200 µm.

**Supplementary Figure 10. Validation of LP_C7 to C9 marker genes with MERFISH data.** Comparison of gene expression profiles for LP cluster 7 to 9 between spatial transcriptomics and publicly available MERFISH data, showing consistent spatial localization. Genes including *MT2b*, *ONI1*, *SCR2*, *ALOG5*, and *TB1*. Scale bars, 200 µm.

**Supplementary Figure 11. Validation of LP_C10 and C11 marker genes with MERFISH data.** Comparison of gene expression profiles for LP cluster 10 and 11 between spatial transcriptomics and publicly available MERFISH data, showing consistent spatial localization. Genes including *GL10*, *RA2*, and *MND1*. Scale bars, 200 µm.

**Supplementary Figure 12. Validation of genes with previously published *in-situ* hybridization results.** Comparison of gene expression profiles for DR and LP between spatial transcriptomics and previously published *in-situ* hybridization data, showing consistent spatial localization. Genes including *ALOG1*, *LAX1/BA1*, and *VRT2*. Scale bars, 200 µm.

**Supplementary Figure 13. Co-expression network in DR stage sections.** Visualization of co-expression modules derived from genes expressed in the DR stage. Modules showed limited separation, likely due to the relatively homogenous gene expression across the tissue. This is consistent with the developmental context, as the DR stage represents an earlier, less differentiated phase of inflorescence development.

**Supplementary Figure 14. Additional gene expression profiles.** Expression patterns of additional genes of interest not shown in main figures.

**Supplementary Figure 15. SEM and paired spikelet distribution in *ps3* mutant compared to WT. a,** Histogram showing the relative frequency of rachis node bins producing paired spikelets in *ps3* spikes (n = 10). Y-axis indicates relative vertical position along the spike (normalized from base [0] to apex [1]). **b**, scanning electron microscopy (SEM) images of WT inflorescence at TS stage. **c-d**, SEM images of *ps3* inflorescences at TS stage with paired spikelet positions highlighted in magenta. Scale bars, 100 μm.

**Supplementary Figure 16. Differentially expressed transcripts (DETs) and GO enrichment in *ps3* RNA-seq. a**, bar plot showing the number of upregulated (cyan) and downregulated (salmon) transcripts in *ps3* relative to WT at three developmental stages: DR, LP, and TS. **b**, Gene Ontology (GO) enrichment analysis of differentially expressed transcripts (DETs). Selected enriched biological processes are shown for each group (DR_DOWN, LP_DOWN, LP_UP, TS_DOWN, TS_UP). Dot size represents gene ratio, and color indicates adjusted p-value.

**Supplementary Figure 17. Histological staining of wheat inflorescence tissue sections used for Stereo-seq.** DR1–DR4 (**a**) and LP1–LP4 (**b**) tissue sections adhered to the surface of the Stereo-seq chip and stained to visualize nuclei and cell walls. Sections were incubated at 37 °C for 2 minutes, fixed in methanol at −20 °C for 30 minutes, and stained with Qubit ssDNA stain (for nuclei; green) and Fluorescent Brightener 28 (for cell walls; blue). Scale bar = 200 μm.

**Supplementary Table 1. Summary of spatial transcriptomic bin statistics for DR and LP samples.** Table includes cell/bin count, bin size, number of detected genes, and summary statistics for molecular identifiers (MID and gene counts) per bin in each sample. The number of bins assigned to each spatial cluster (1–11) is also provided for DR (bin40) and LP (bin50) stages.

**Supplementary Table 2. Marker genes for DR and LP spatial clusters.** List of marker genes identified across spatial clusters in DR and LP stages.

**Supplementary Table 3. Cluster-enriched genes.** Genes significantly enriched in individual spatial clusters based on expression patterns.

**Supplementary Table 4. Gene lists from pseudotime analysis.** Full list of genes assigned to pseudotime expression profiles in DR and LP trajectories.

**Supplementary Table 5. Gene modules from LP co-expression analysis.** Gene list for modules identified through co-expression network analysis in LP stage.

**Supplementary Table 6. Transcript abundance of differentially expressed transcripts in *ps3* relative to WT.** TPM (Transcripts Per Million) values for differentially expressed transcripts at the DR, LP and TS stages, comparing *ps3* and wild-type. Data are provided separately for transcripts upregulated or downregulated at each stage and represent the mean of three biological replicates.

**Supplementary Table 7. Transcript quantities of leaf–ridge/spikelet boundary genes in WT and *ps3*.** TPM values for selected leaf-ridge/spikelet boundary-associated genes measured at double ridge (DR), lemma primordium (LP), and terminal spikelet (TS) stages in WT and *ps3*. Data represent the mean of three biological replicates. Fold-change values (*ps3*/WT) for each stage are also included.

