## Supplementary figures for "Spatial transcriptomics identifies distinct domains regulating yield-related traits of the wheat ear"

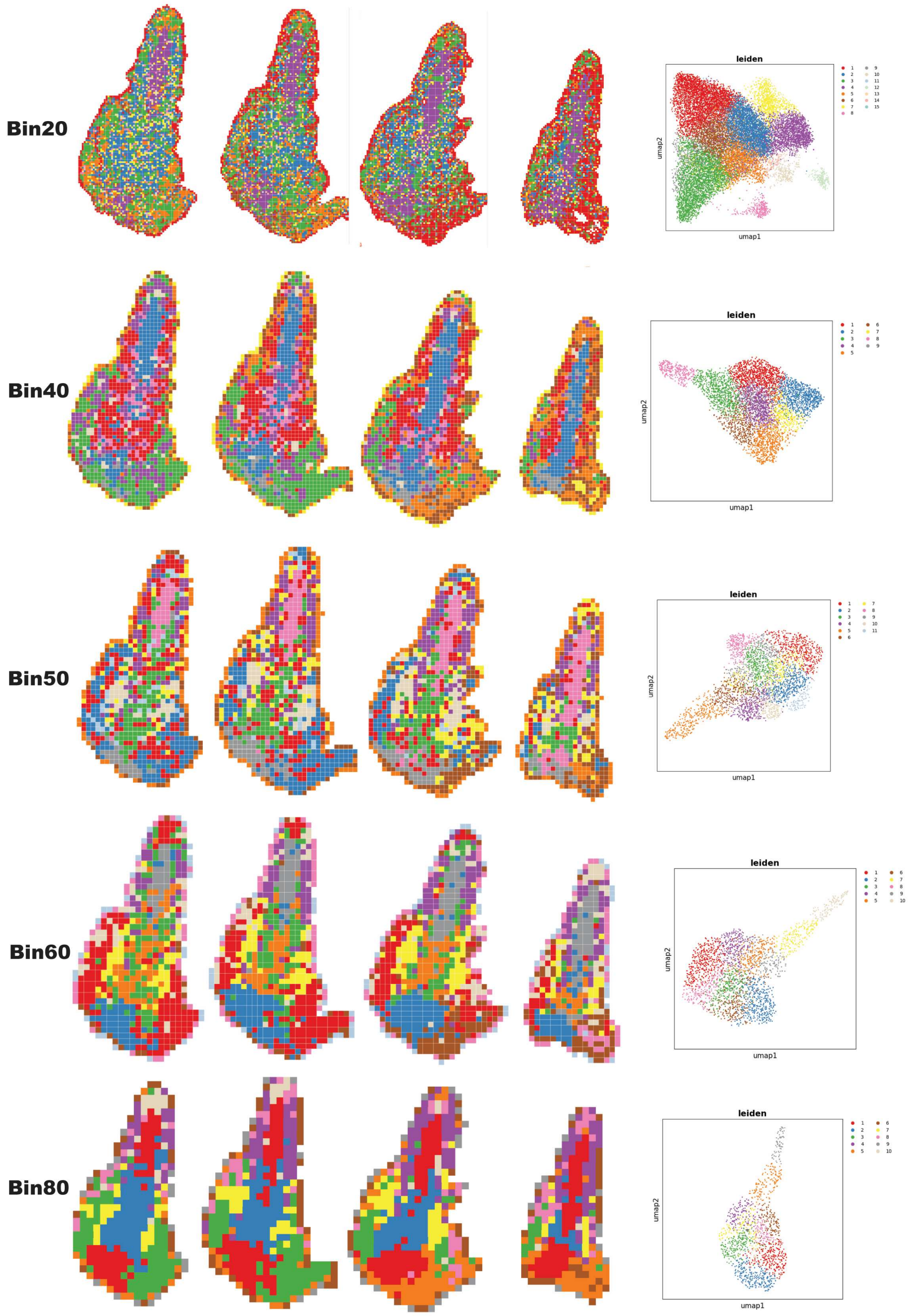


**Supplementary Figure 1. Overview of spatial resolution at different bin sizes in double ridge (DR) stage sections.** Spatial maps and UMAP projections of DR stage sections showing cluster resolution at bin sizes including Bin20, Bin40, Bin50, Bin60, and Bin80.

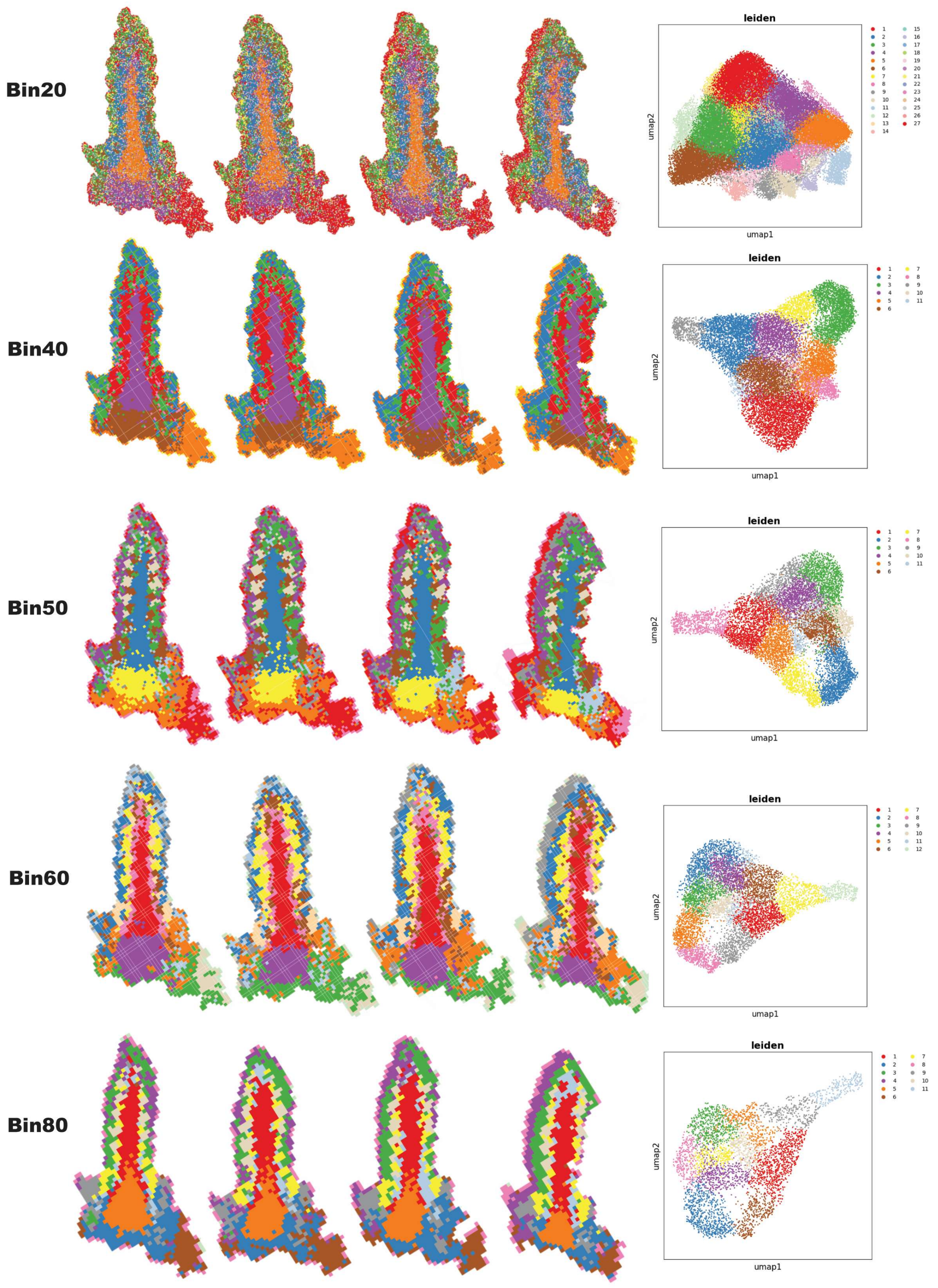


**Supplementary Figure 2. Overview of spatial resolution at different bin sizes in lemma primordia (LP) stage sections.** Spatial maps and UMAP projections of LP stage sections showing cluster resolution at bin sizes including Bin20, Bin40, Bin50, Bin60, and Bin80.

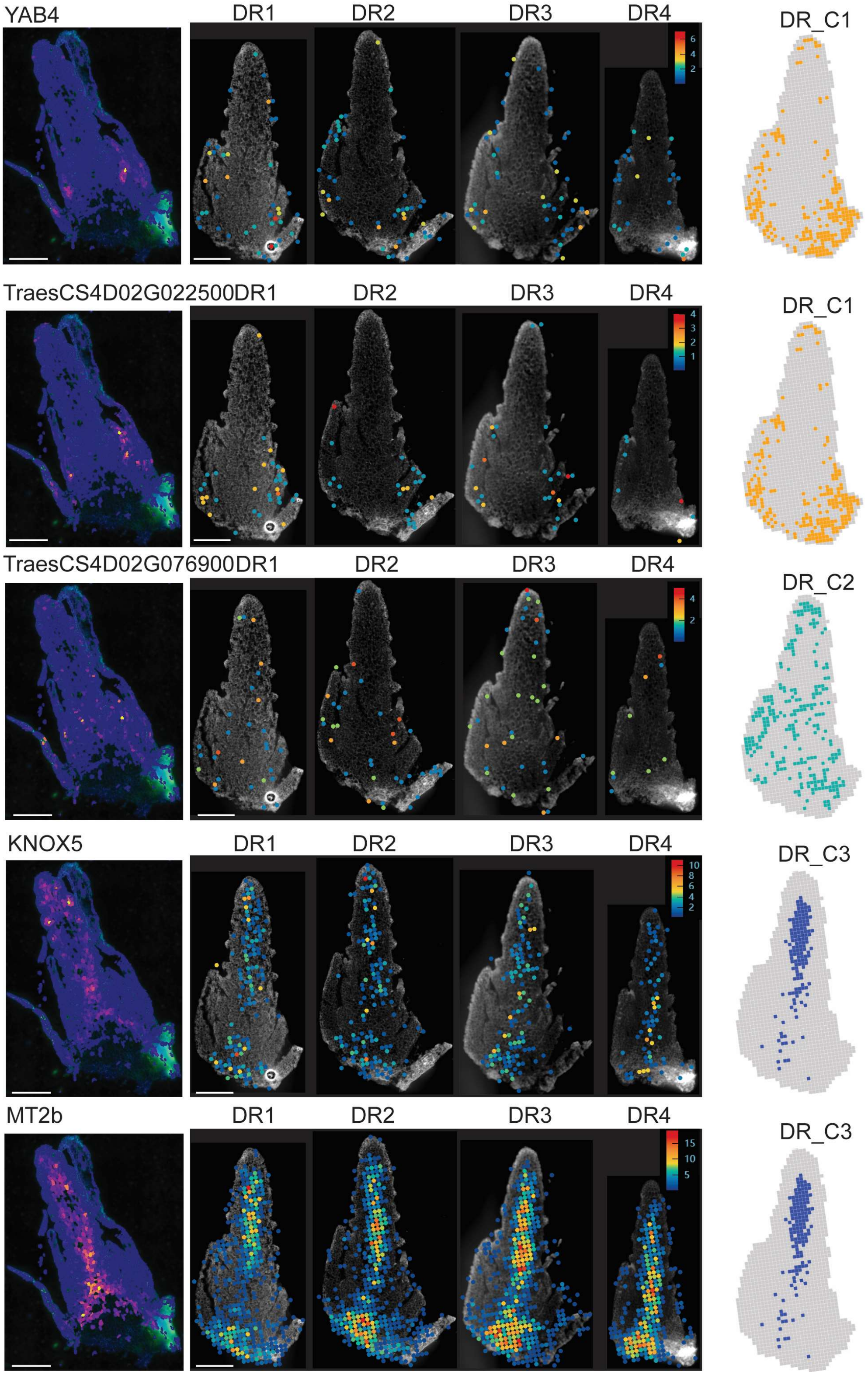


**Supplementary Figure 3. Validation of DR_C1 to C3 marker genes with MERFISH data.** Comparison of gene expression profiles for DR cluster 1 to 3 between spatial transcriptomics and publicly available MERFISH data[1], showing consistent spatial localization. Genes including *YAB4*, *TraesCS4D02G022500*, *TraesCS4D02G076900*, *KNOX5*, and *MT2b*. Scale bars, 200 μm.

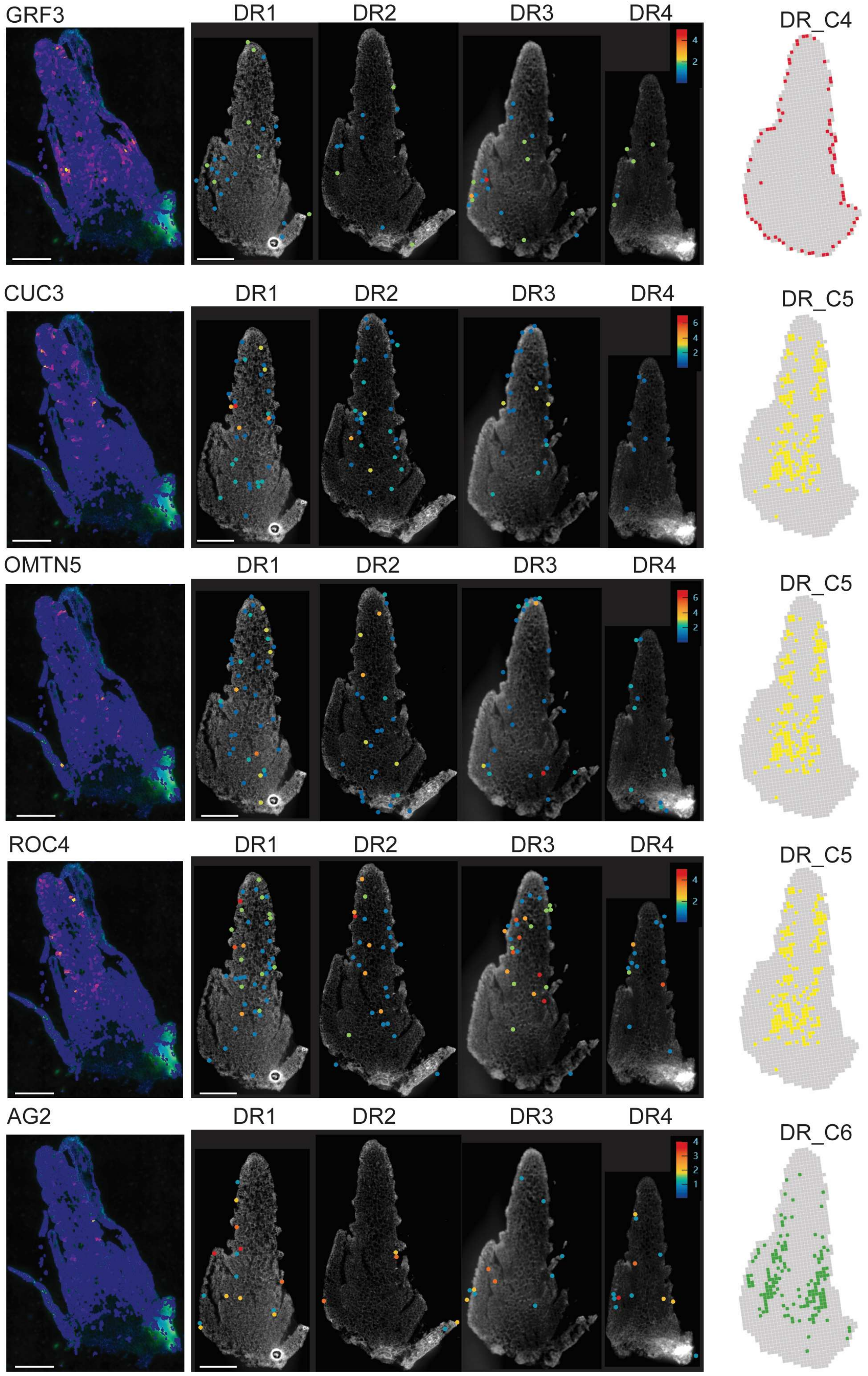


**Supplementary Figure 4. Validation of DR_C4 to C6 marker genes with MERFISH data.** Comparison of gene expression profiles for DR cluster 4 to 6 between spatial transcriptomics and publicly available MERFISH data[1], showing consistent spatial localization. Genes including *GRF3*, *CUC3*, *OMTN5*, *ROC4*, and *AG2*. Scale bars, 200 μm.

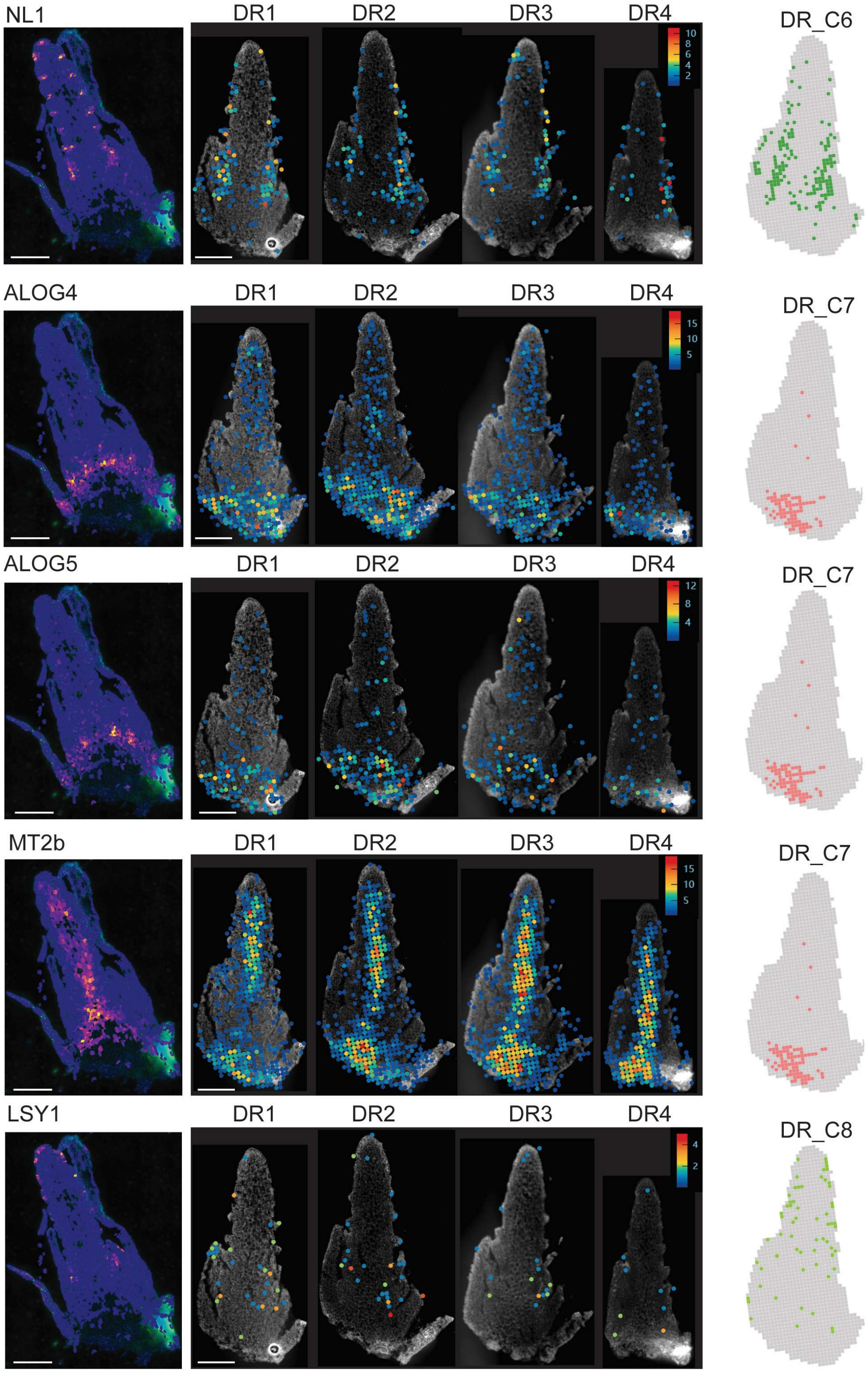


**Supplementary Figure 5. Validation of DR_C6 to C8 marker genes with MERFISH data.** Comparison of gene expression profiles for DR cluster 6 to 8 between spatial transcriptomics and publicly available MERFISH data[1], showing consistent spatial localization. Genes including *NL1*, *ALOG4*, *ALOG5*, *MT2b*, and *LSY1*. Scale bars, 200 μm.

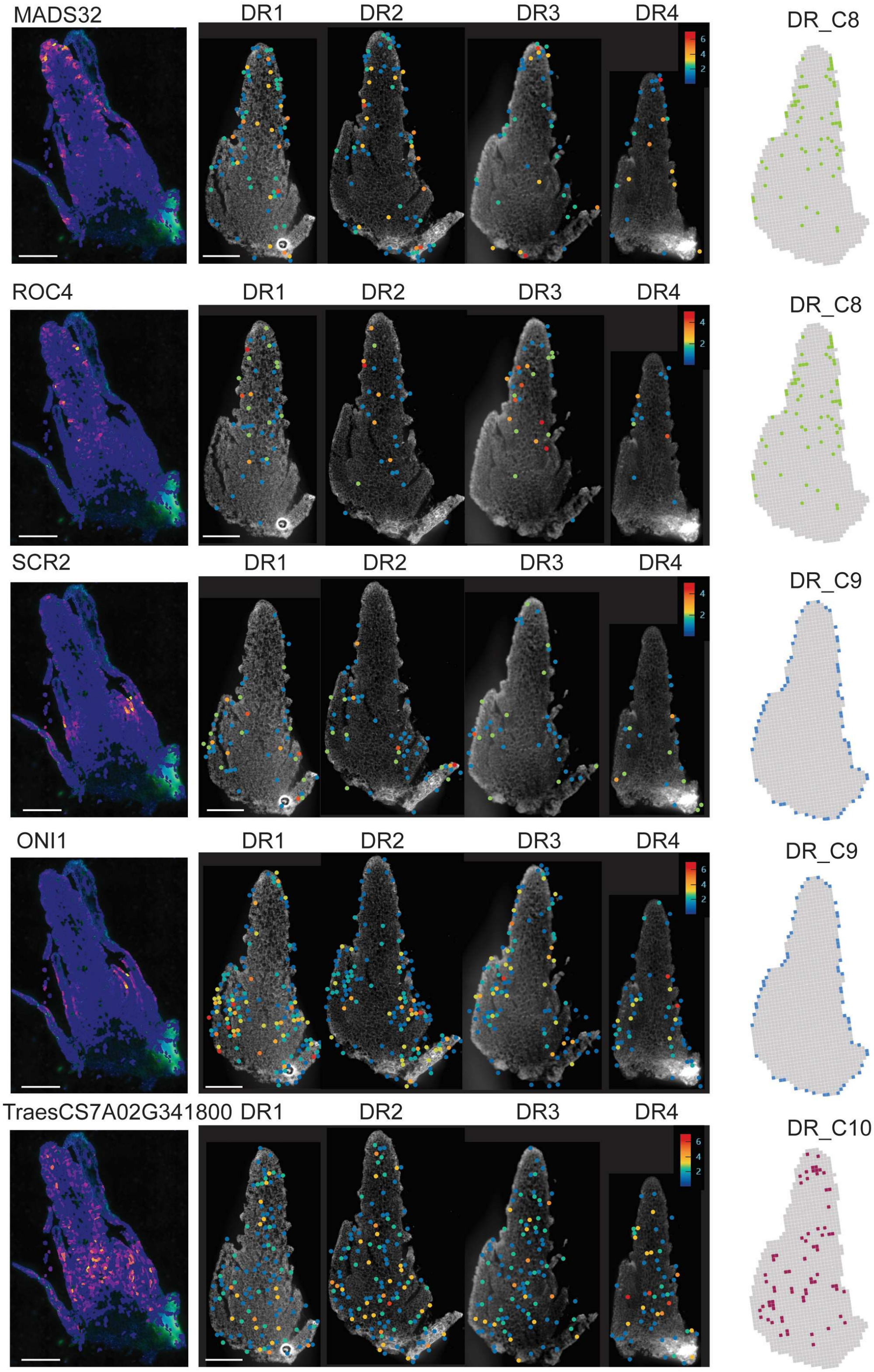


**Supplementary Figure 6. Validation of DR_C8 to C10 marker genes with MERFISH data.** Comparison of gene expression profiles for DR cluster 8 to 10 between spatial transcriptomics and publicly available MERFISH data[1], showing consistent spatial localization. Genes including *MADS32*, *ROC4*, *SCR2*, *ONI1*, and *TraesCS7A02G341800*. Scale bars, 200 μm.

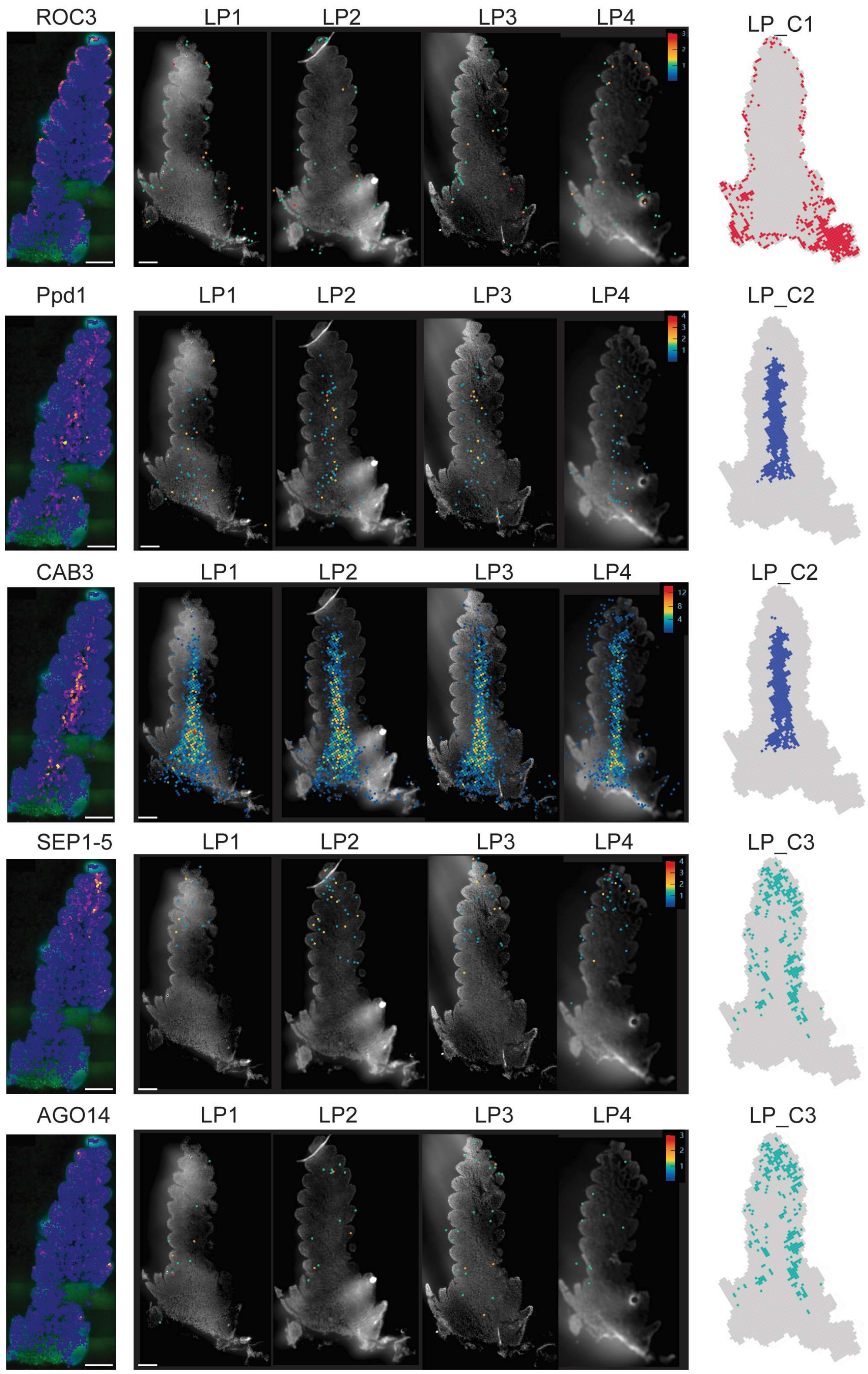


**Supplementary Figure 7. Validation of LP_C1 to C3 marker genes with MERFISH data.** Comparison of gene expression profiles for LP cluster 1 to 3 between spatial transcriptomics and publicly available MERFISH data[1], showing consistent spatial localization. Genes including *ROC3*, *Ppd1*, *CAB3*, *SEP1-5*, and *AGO14*. Scale bars, 200 μm.

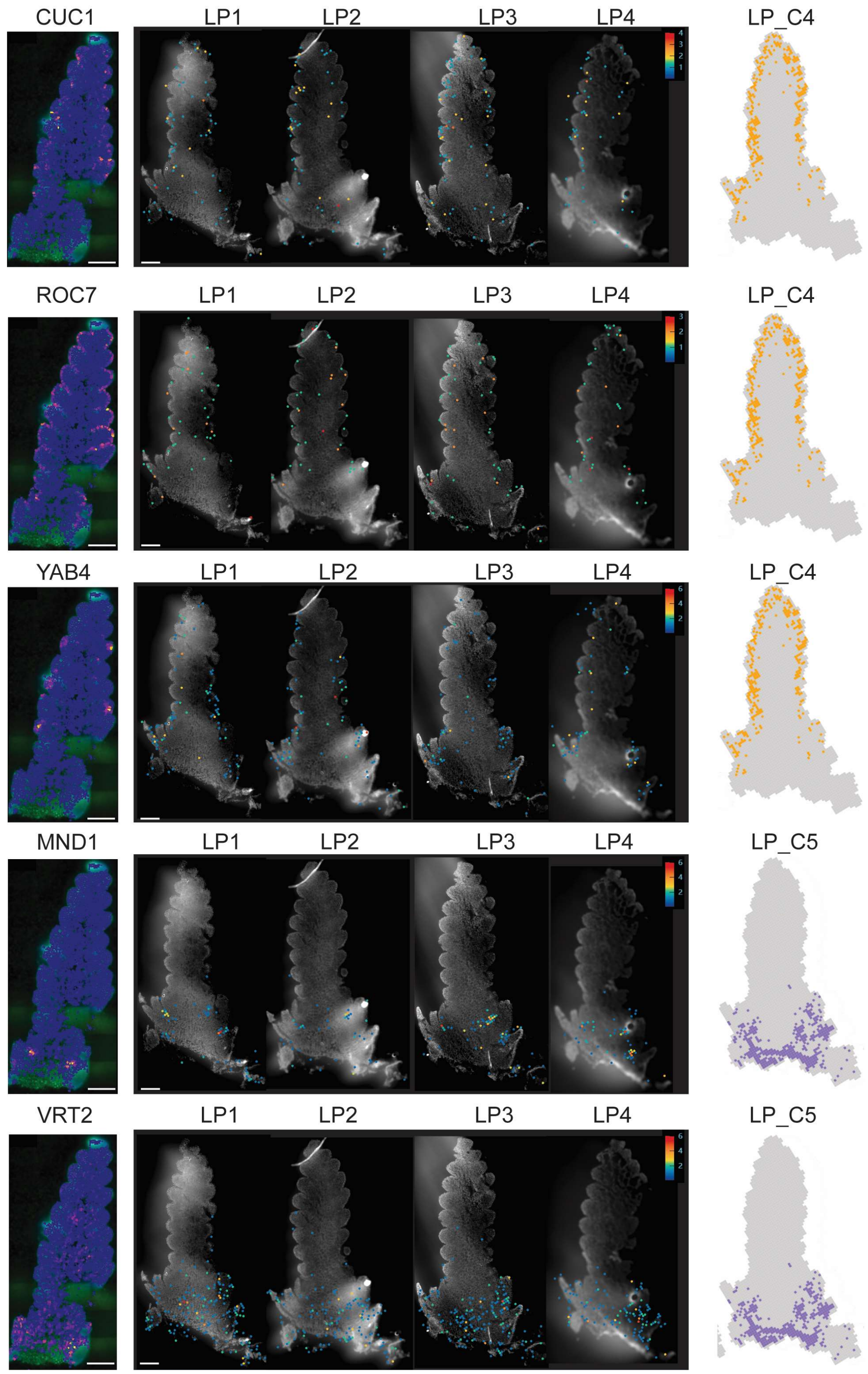


**Supplementary Figure 8. Validation of LP_C4 and C5 marker genes with MERFISH data.** Comparison of gene expression profiles for LP cluster 4 and 5 between spatial transcriptomics and publicly available MERFISH data[1], showing consistent spatial localization. Genes including *CUC1*, *ROC7*, *YAB4*, *MND1*, and *VRT2*. Scale bars, 200 μm.

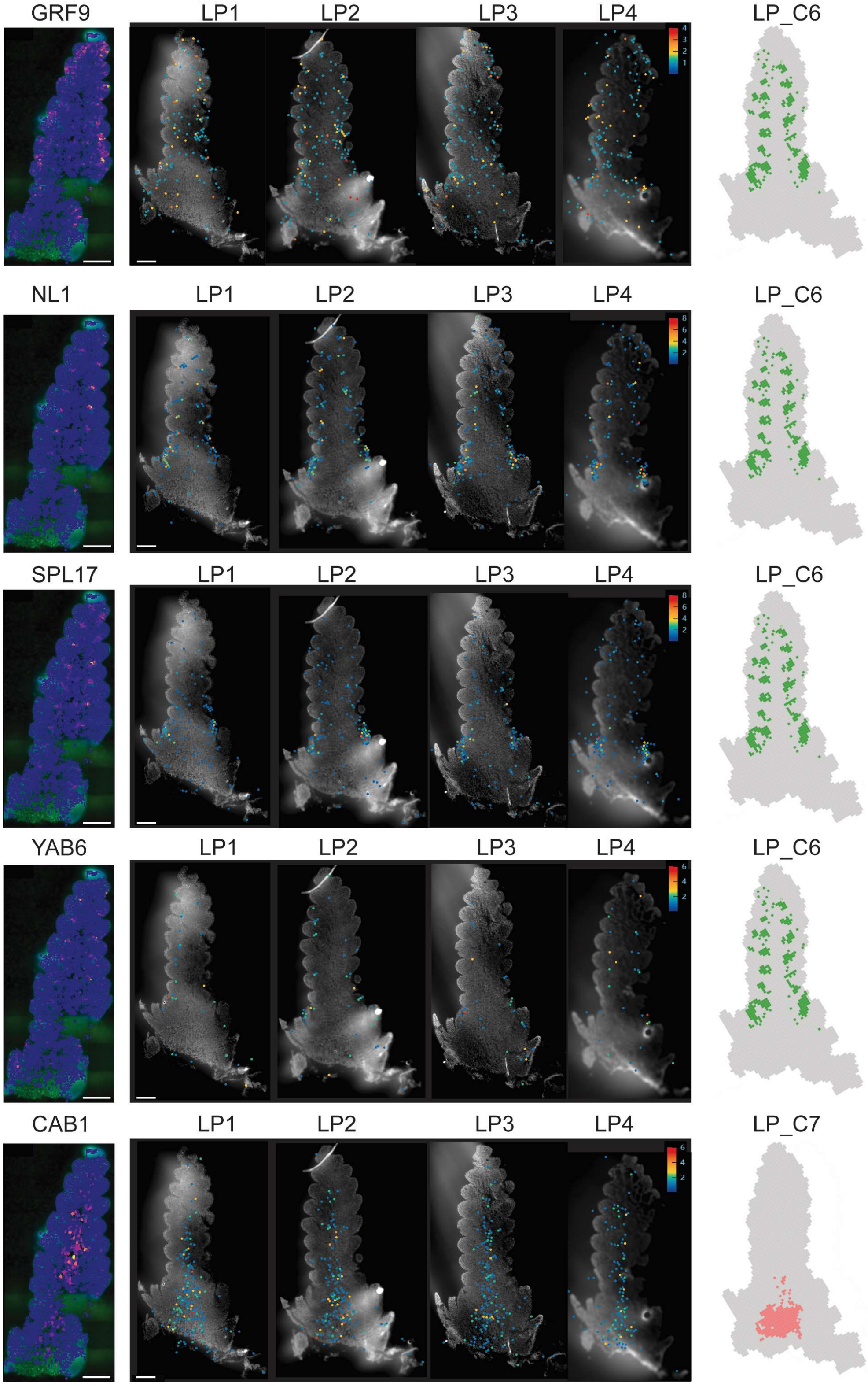


**Supplementary Figure 9. Validation of LP_C6 and C7 marker genes with MERFISH data.** Comparison of gene expression profiles for LP cluster 6 and 7 between spatial transcriptomics and publicly available MERFISH data[1], showing consistent spatial localization. Genes including *GRF9*, *NL1*, *SPL17*, *YAB6*, and *CAB1*. Scale bars, 200 μm.

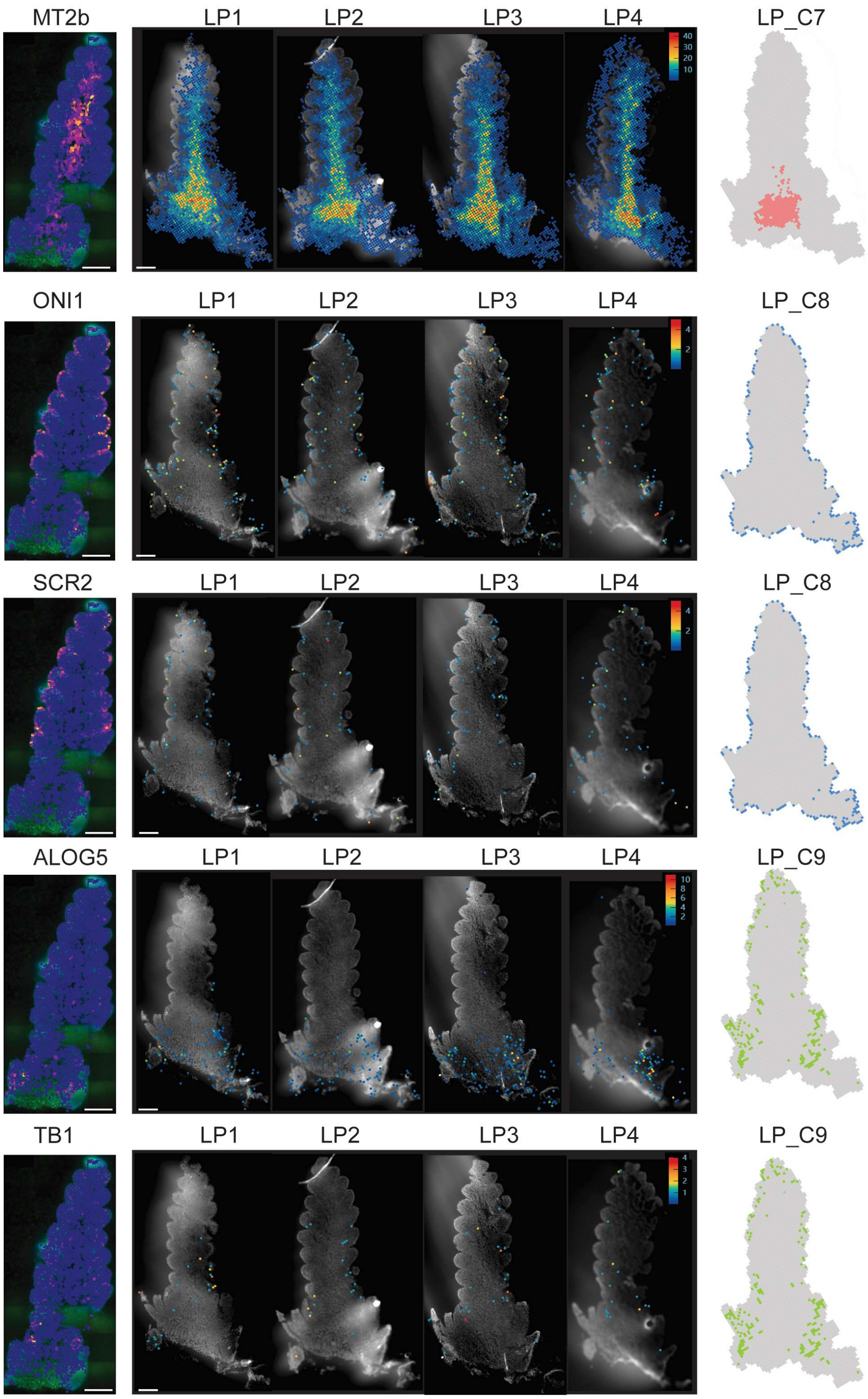


**Supplementary Figure 10. Validation of LP_C7 to C9 marker genes with MERFISH data.** Comparison of gene expression profiles for LP cluster 7 to 9 between spatial transcriptomics and publicly available MERFISH data[1], showing consistent spatial localization. Genes including *MT2b*, *ONI1*, *SCR2*, *ALOG5*, and *TB1*. Scale bars, 200 μm.

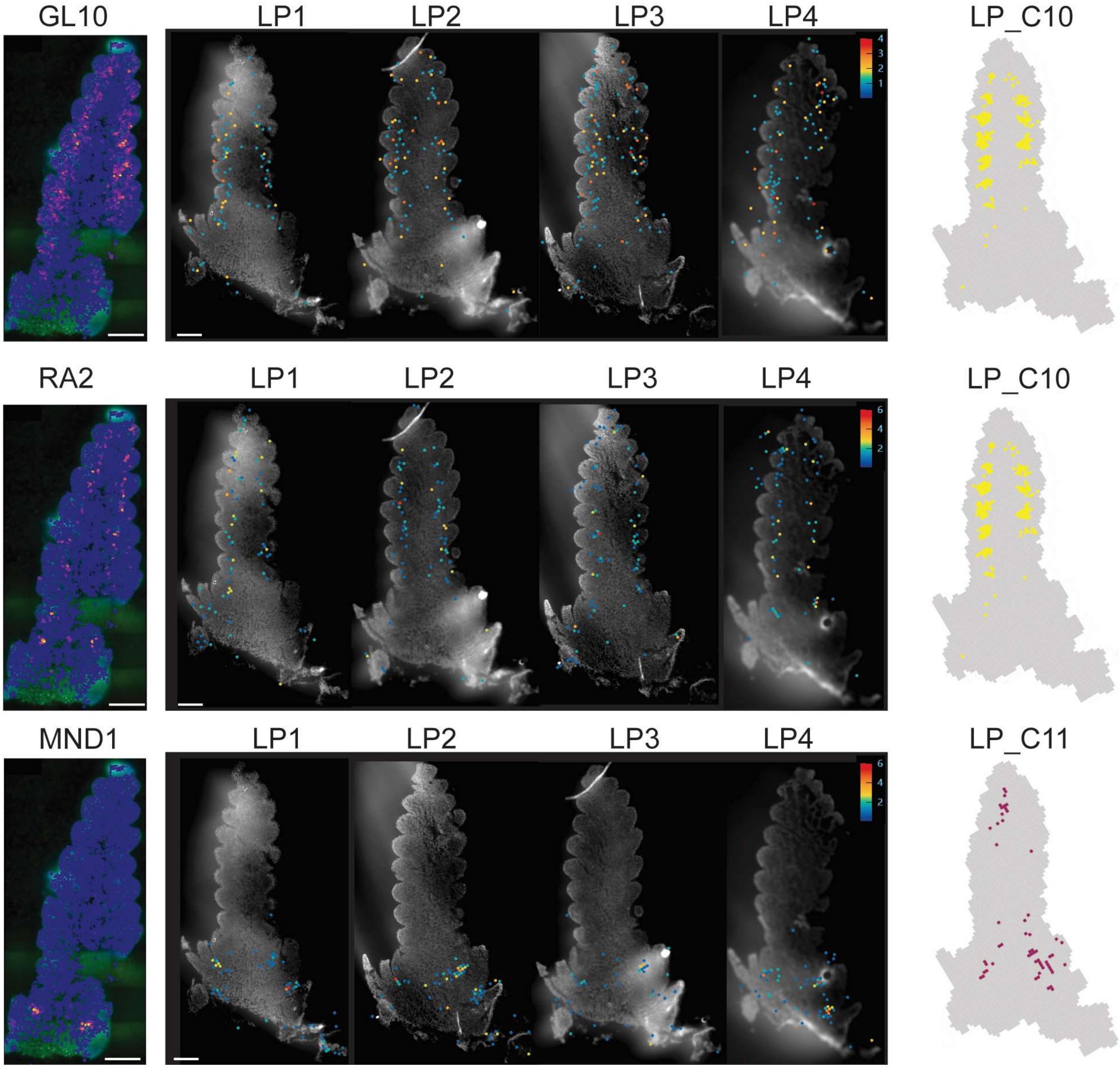


**Supplementary Figure 11. Validation of LP_C10 and C11 marker genes with MERFISH data.** Comparison of gene expression profiles for LP cluster 10 and 11 between spatial transcriptomics and publicly available MERFISH data[1], showing consistent spatial localization. Genes including *GL10*, *RA2*, and *MND1*. Scale bars, 200 µm.

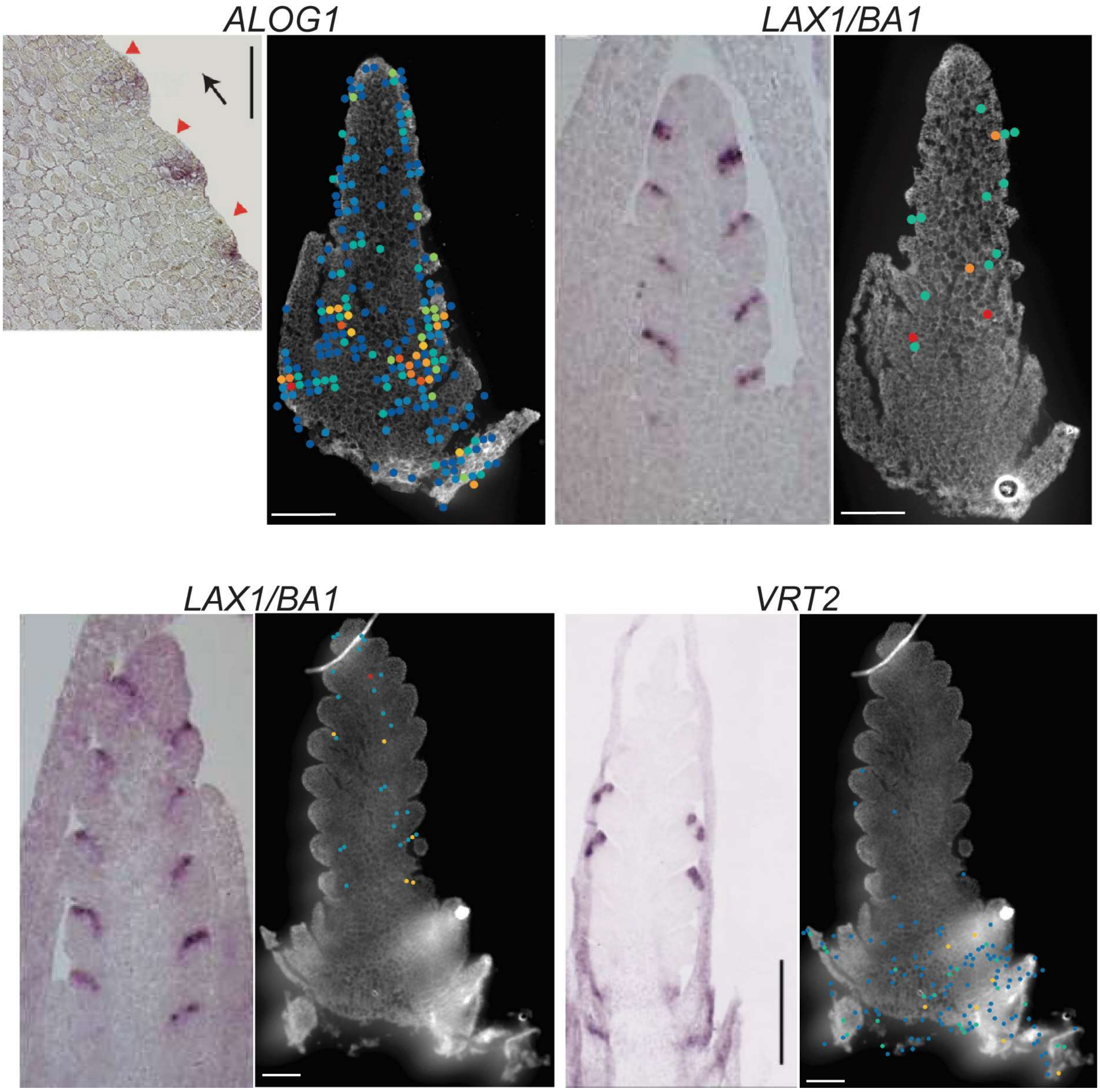


**Supplementary Figure 12. Validation of genes with previously published *in-situ* hybridization results.** Comparison of gene expression profiles for DR and LP between spatial transcriptomics and previously published *in-situ* hybridization data, showing consistent spatial localization. Genes including *ALOG1*[2], *LAX1/BA1*[3], and *VRT2*[4]. Scale bars, 200 μm.

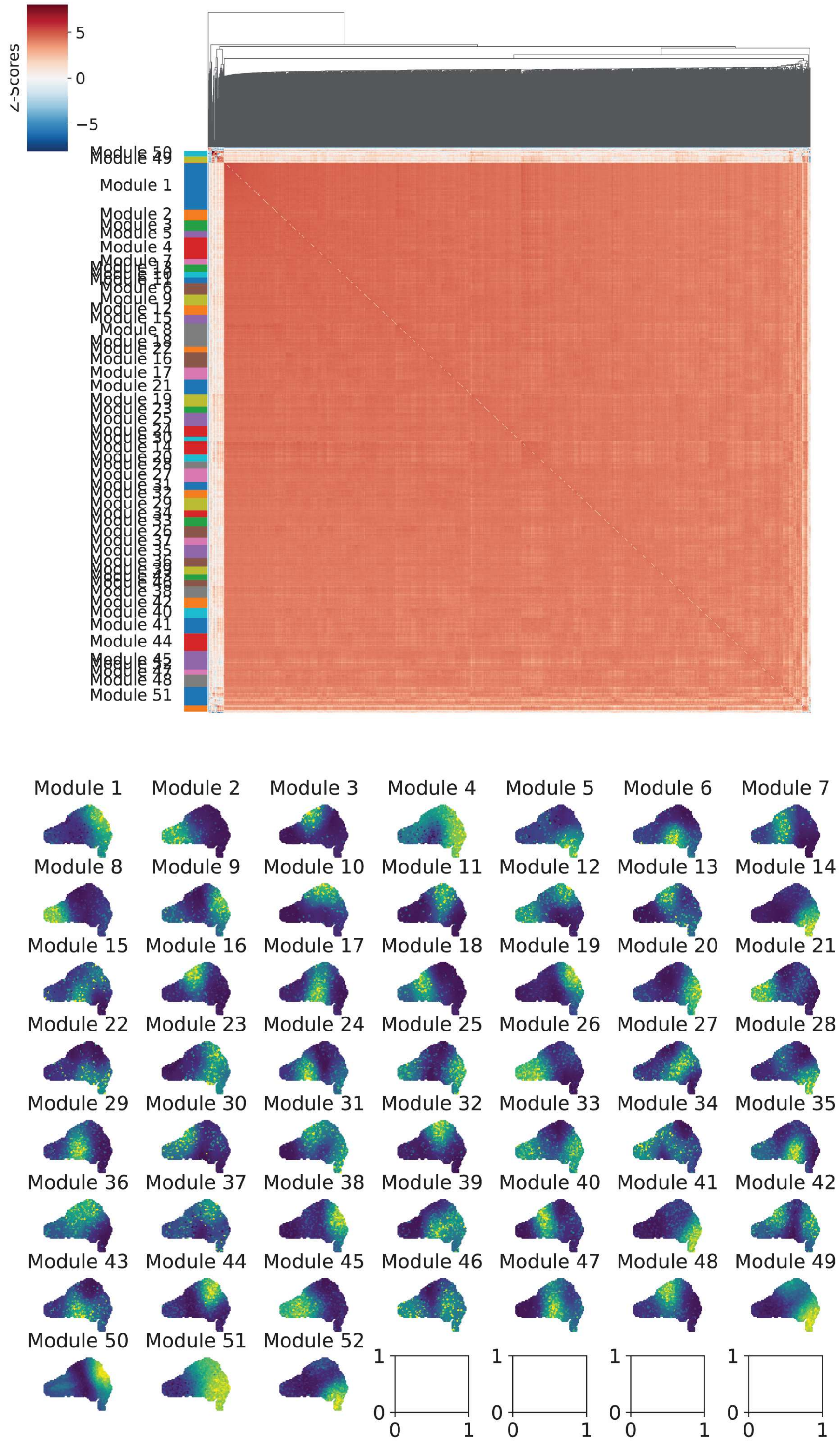


**Supplementary Figure 13. Co-expression network in DR stage sections.** Visualization of co-expression modules derived from genes expressed in the DR stage. Modules showed limited separation, likely due to the relatively homogenous gene expression across the tissue. This is consistent with the developmental context, as the DR stage represents an earlier, less differentiated phase of inflorescence development.

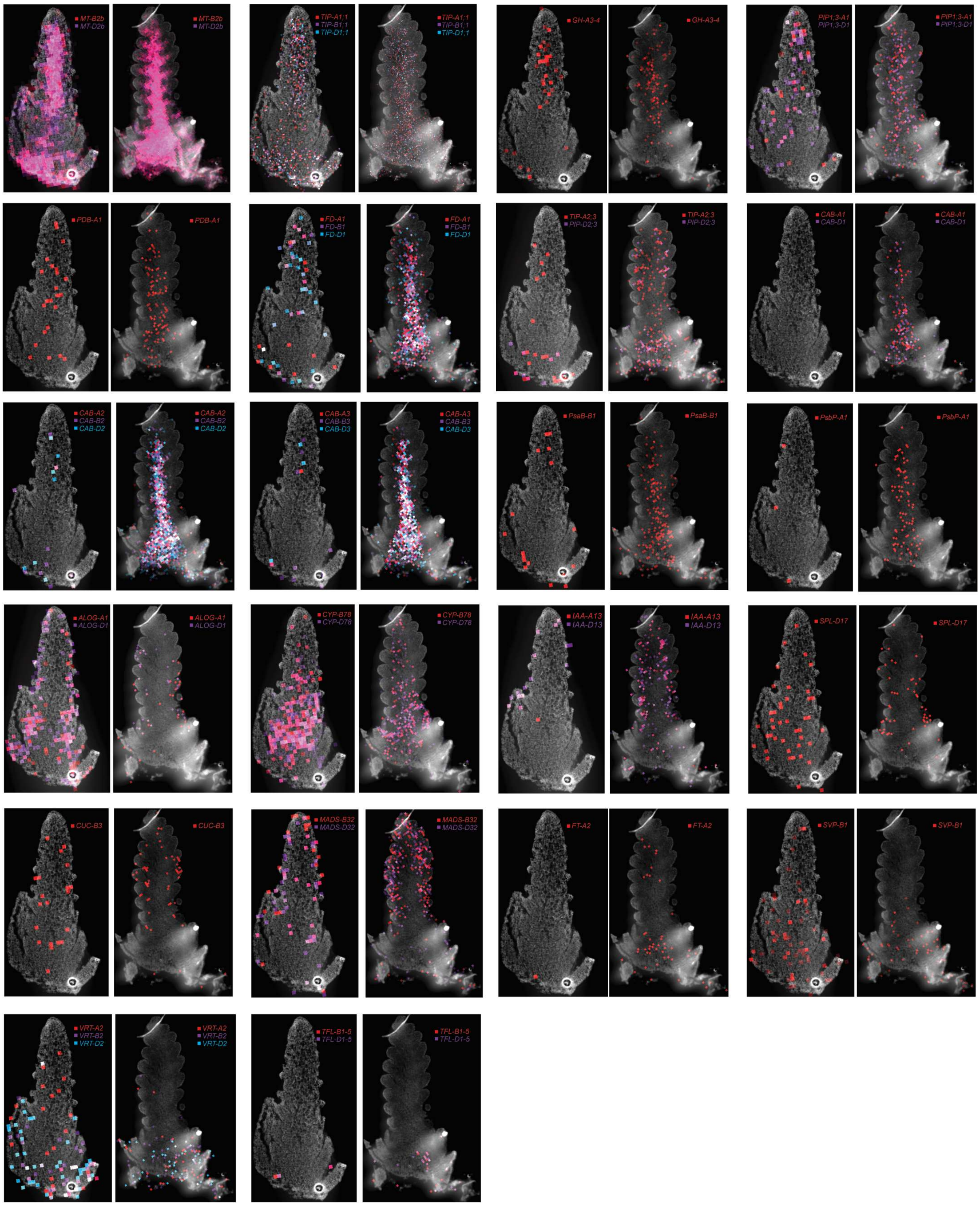


**Supplementary Figure 14. Additional gene expression profiles.** Expression patterns of additional genes of interest not shown in main figures.

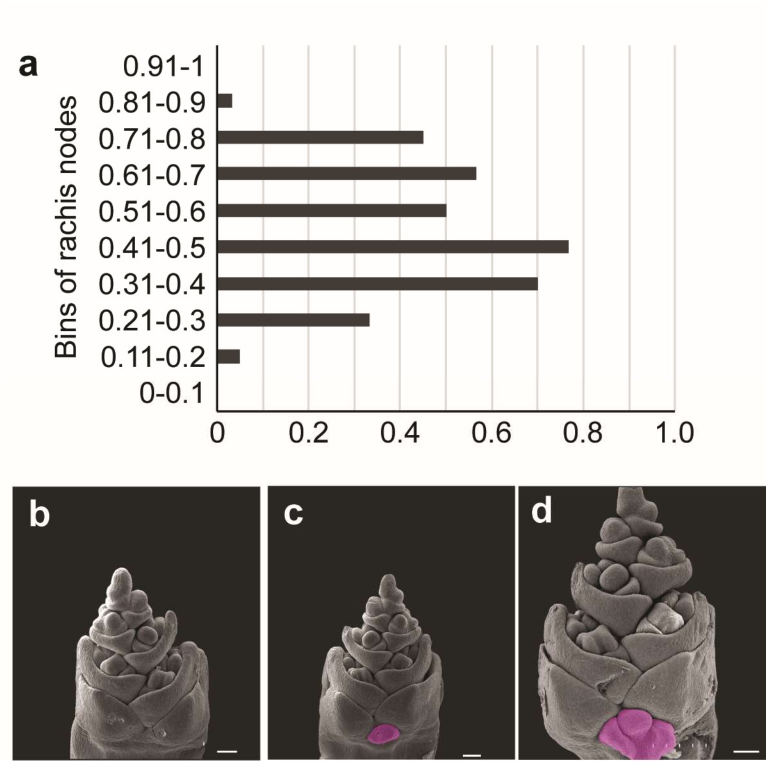


**Supplementary Figure 15. SEM and paired spikelet distribution in *ps3* mutant compared to WT. a,** Histogram showing the relative frequency of rachis node bins producing paired spikelets in *ps3* spikes (n = 10). Y-axis indicates relative vertical position along the spike (normalized from base [0] to apex [1]). **b**, scanning electron microscopy (SEM) images of WT inflorescence. **c-d**, SEM images of *ps3* inflorescences with paired spikelet positions highlighted in magenta. Scale bars, 100 μm.

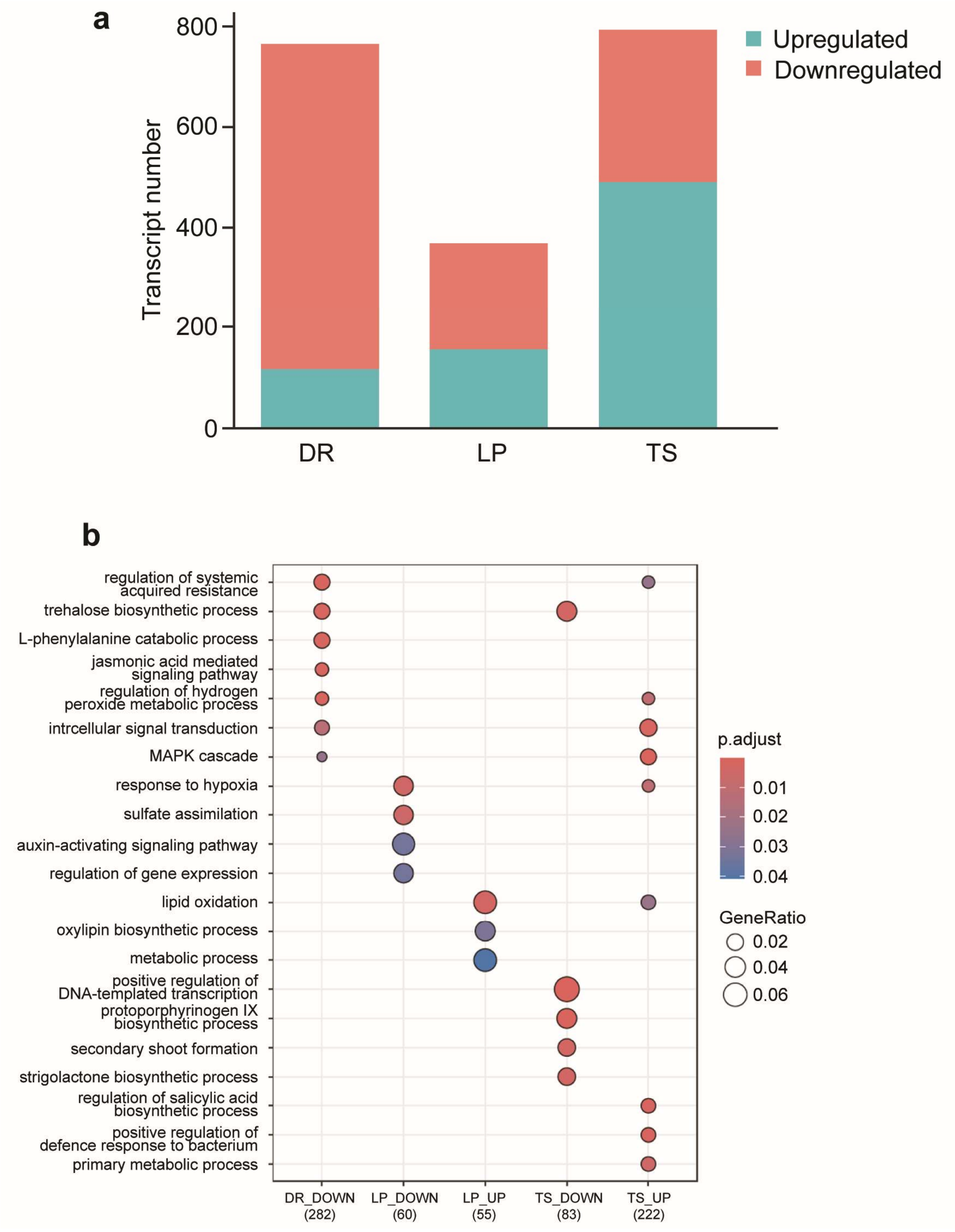


**Supplementary Figure 16. Differentially expressed transcripts (DETs) and GO enrichment in *ps3* RNA-seq. a**, bar plot showing the number of upregulated (cyan) and downregulated (salmon) transcripts in *ps3* relative to WT at three developmental stages: DR, LP, and TS. **b**, Gene Ontology (GO) enrichment analysis of differentially expressed transcripts (DETs). Selected enriched biological processes are shown for each group (DR_DOWN, LP_DOWN, LP_UP, TS_DOWN, TS_UP). Dot size represents gene ratio, and color indicates adjusted p-value.

a

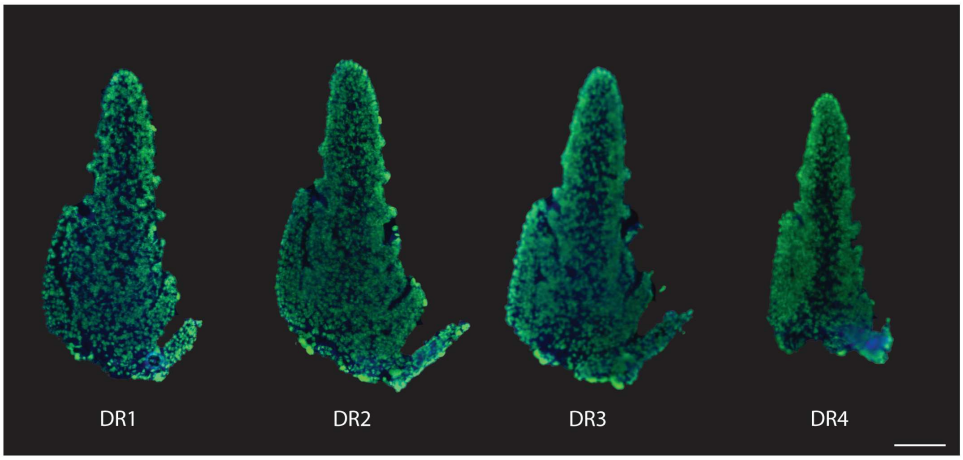


b

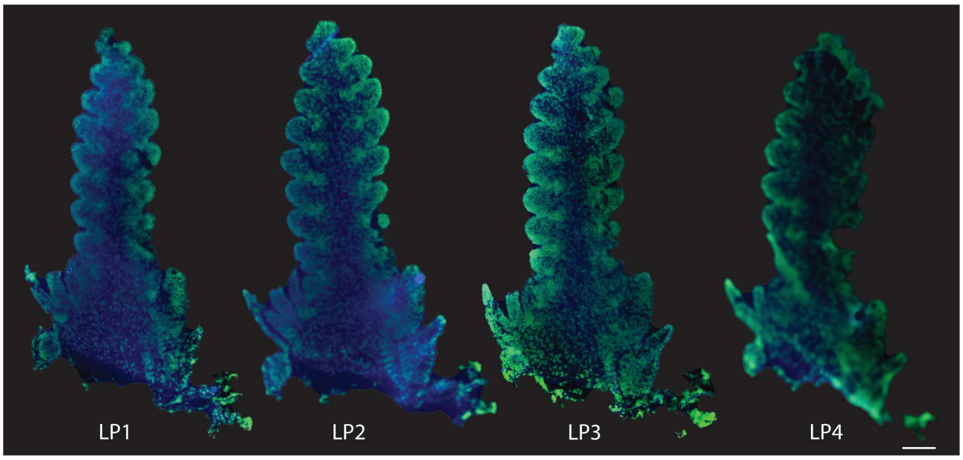


**Supplementary Figure 17. Histological staining of wheat inflorescence tissue sections used for Stereo-seq.** DR1–DR4 (**a**) and LP1–LP4 (**b**) tissue sections adhered to the surface of the Stereo-seq chip and stained to visualize nuclei and cell walls. Sections were incubated at 37 °C for 2 minutes, fixed in methanol at −20 °C for 30 minutes, and stained with Qubit ssDNA stain (for nuclei; green) and Fluorescent Brightener 28 (for cell walls; blue). Scale bar = 200 μm.
